# Lack of Purple Acid Phosphatase *SlPAP26b* compromises the phosphorus starvation response in tomato independent of SlPHR1 and SlPHL1

**DOI:** 10.1101/2023.11.29.569173

**Authors:** Akash, Rajat Srivastava, Abhishek Roychowdhury, Kapil Sharma, Martin Cerny, Pavel Kerchev, Rahul Kumar

**Author notes:** Equal contribution (Akash, Rajat Srivastava). Corresponding address Dr. Rahul Kumar Department of Plant Sciences, School of Life Sciences, University of Hyderabad, Hyderabad, 500046, Telangana, India.

## Abstract

The scarcity of soil phosphorus (P), an essential macronutrient, often limits plant growth and development. Enhanced secretion of intracellular and secretory acid phosphatases is essential to maintain cellular inorganic P (Pi) homeostasis in plants. Herein, using transcriptomics and proteomics approach, we observed upregulation of several purple acid phosphatases (PAPs), including *SlPAP1*, *SlPAP10b*, *SlPAP12*, *SlPAP15*, *SlPAP17b*, *SlPAP26a*, and *SlPAP26b* in Pi-deficient tomato seedlings. Higher transcript levels of *SlPAP17b* and *SlPAP26b* in the older senescing leaves than the younger leaves indicated active involvement of these PAPs in Pi remobilization. Subsequent detailed characterization of *SlPAP17b*, *SlPAP26a*, and *SlPAP26b* revealed a prominent role of *SlPAP26b* in Pi homeostasis. Silencing of *SlPAP26b* led to an exacerbated P starvation response as these plants exhibited smaller shoots, lower soluble Pi, total P levels, and higher sucrose than their EV controls under Pi deprivation. *SlPAP26b-*silenced plants also showed misregulation of P starvation inducible genes such as phosphate transporters and glycerolipid remodellers, even under Pi-sufficient conditions. Whereas *SlPAP26b* levels were induced by external sucrose, its expression was found to be independent of the Myb class master regulators of P starvation response, SlPHR1 and SlPHL1. Altogether, this study identifies a prominent role of *SlPAP26b* in the Pi compensation network in tomato seedlings.

## Introduction

Phosphorus (P), an essential macronutrient, is critical for plant growth and development. P is a necessary constituent of essential biomolecules such as nucleic acids, phospholipids, cellular energy currency, and ATP (Roychowdhury et al., 2023). It also participates in protein phosphorylation and cell signaling events. In soil roots assimilate P in the form of inorganic phosphate (Pi), predominantly available in three orthophosphate species (H_3_PO_4_, H_2_PO_4_^−^, and HPO_4_^2–^) (Schachtman et al., 1998). Due to its poor mobility in the soil solution, Pi often becomes a limiting nutrient in agriculture and the application of large quantities of phosphate fertilizers is practiced to mitigate its deficiency (Hammond et al., 2003). However, owing to its high reactivity and rapid fixation in soils, the nutrient-use-efficiency of the conventional Pi-fertilizers is often poor, even under a well-designed Pi fertilization scheme (Syers et al., 2008). A major fraction of soil P (up to 80%) remains in the form of organic compounds, including phosphate monoesters and phosphate diesters (nucleic acids, phospholipids, cyclic phosphates) and phosphonates (George and Richardson, 2008). Pi also reacts with cations such as aluminum, iron or calcium, and magnesium in a pH-dependent manner to form insoluble complexes in soils (Jez et al., 2016; Péret et al., 2011). Thus, despite the availability of a significant amount of P in its organic forms or insoluble mineral-P complexes, plants often experience Pi deficiency. Plants employ various morphological, physiological, biochemical, and molecular adaptive measures to cope with Pi shortage. Some of these responses include reprogramming root system architecture (RSA), carbon repartitioning between shoot and root, enhanced secretion of organic acids and carboxylates, and association with mycorrhizal fungi. Transcription, activity, and exudation of ribonucleases and acid phosphatases also increase under Pi deprivation (Plaxton and Tran, 2011; Secco et al., 2013; Srivastava et al., 2021a).

Up-regulation of intracellular acid phosphatase (IAP) and secreted acid phosphatase (SAP) enzymes (APases; E.C. 3.1.3.2), involved in the hydrolysis of Pi from various phosphate monoesters and anhydrides in the acidic pH range, is a well-defined Pi starvation response exhibited by plants (Tran et al., 2010). The enhanced APase activity in Pi-deficient plants is contributed by the induction of several purple acid phosphatases (González-Muñoz et al., 2015; Li et al., 2002; Mehra et al., 2017; Srivastava et al., 2020). While intracellular acid phosphatases are involved in recycling Pi from dispensable intracellular P-pools, secreted acid phosphatases act upon external phosphoesters or inorganic mineral-P complexes to mobilize Pi in the apoplast or rhizosphere (Bozzo and Plaxton, 2008; Vance et al., 2003). Purple acid phosphatases (PAPs), the predominant class of APases, are present in diverse organisms including bacteria, fungi, and plants (Kuang et al., 2009). PAPs belong to a large multigene family of plants. The full complement of PAPs has been identified in many species, including arabidopsis (Li et al., 2002), rice (Zhang et al., 2011), soybean (Li et al., 2012), maize (González-Muñoz et al., 2015) and tomato (Srivastava et al., 2020).

In Arabidopsis, *AtPAP17* and *AtPAP26* are involved in Pi remobilization and Pi homeostasis. Upregulation of *PAP26*, a dual-targeted PAP with vacuolar acid phosphatase and secreted APase activity, regulates Pi remobilization, ATP utilization, and senescence in Arabidopsis seedlings (Hurley et al., 2010; Robinson et al., 2012). *AtPAP15*, a purple acid phosphatase with phytase activity, influences seed and pollen germination in Arabidopsis (Kuang et al., 2009). Several PAPs have also been characterized for their roles in phosphate starvation response (PSR) in rice seedlings (Farhadi et al., 2020). For example, *OsPAP21b* hydrolyzes the P monoester and anhydride from extracellular and intracellular P pools to release Pi for its use in metabolic processes (Mehra et al., 2017). A root-secreted *OsPAP3b* has been recently reported to utilize organic phosphate and promote plant growth in the transgenic lines overexpressing this gene (Bhadouria et al., 2023). *OsPAP26* has been reported to regulate phosphate remobilization in rice (Gao et al. 2017). In soybean, *GmPAP4* overexpression results in improved P use efficiency. *GmPAP14* influences the utilization of phytate for improved plant growth and development. PAP10 orthologs in plants have been reported to be primarily root-specific with a major role in Pi uptake (Lu et al., 2016; Tian et al., 2012).

Myb transcription factors, PHR1 (PHOSPHATE STARVATION RESPONSE1), and PHR1-Like (PHL) orthologs in plants are primary regulators of the transcriptional activity of P starvation inducible genes (Rubio et al., 2001). To date, many direct targets of PHRs such as phosphate transporters, glycerophosphodiestereases (GDPDs), non-protein-coding transcript INDUCED BY PHOSPHATE STARVATION1 (IPS1), miR399, and acid phosphatases have been identified (Bustos et al., 2010; Mehra and Giri, 2016). The transcriptional activity of PHR1 orthologs is controlled by SPX (named after <u>S</u>YG1, <u>P</u>HO81, and <u>X</u>pr1 of yeast) proteins, the negative regulators of Pi signaling. Members of the SPX gene family physically interact with PHR1 and its orthologs and block their binding to the PHOSPHATE STARVATION RESPONSE1 binding site (P1BS, GNATATNC) of the target phosphate starvation inducible (PSI) genes to control P starvation response under non-limiting Pi conditions (Chien et al., 2018; Puga et al., 2014; Wang et al., 2014a). Under Pi starvation, SPX proteins undergo proteasomal degradation, releasing PHRs to bind to PSI gene promoters and initiate P starvation response (Bustos et al., 2010; Rubio et al., 2001). Besides their tight regulation by the SPX-PHR1 module, PSI genes expression is also influenced by other transcription factors and plant carbohydrate status. Accumulated evidence suggests the potential role of shoot-derived sugar signals in initiating acclamatory responses in plant roots under Pi-limiting conditions. These shoot-derived sugar molecules are known to act as systemic plant growth regulators (Hammond and White, 2008). The enhanced shoot-to-root sugar translocation is important for stronger activation of several PSI genes, elevated secretory acid phosphatase activity, and organic acid secretion (Abdelrahman et al., 2018; Akash et al., 2021; Hammond and White, 2008).

In tomato, three PSI acid phosphatases with distinctive physical and kinetic properties have been identified previously (G. Bozzo et al., 2004; G. Bozzo 2002). A PSI *LePS2* gene has also been reported to exhibit acid phosphatase activity (Baldwin et al., 2001). Recently, we reported the presence of 25 PAPs in tomato genome and identified two isoforms each of SlPAP17 (*SlPAP17a/SlPAP17b*) and SlPAP26 (*SlPAP26a*/*SlPAP26b*) (Srivastava et al., 2020). However, no detailed study of any tomato PAP has been carried out for their functions using tomato as a system. Further, studies encompassing their transcriptional regulation and loss of function are missing. In the present study, we profiled the complete set of PSI PAPs using transcriptome and proteome analyses. Among the commonly identified members in both datasets, *SlPAP17b*, *SlPAP26a,* and *SlPAP26b* were selected for their detailed characterization. Functional characterization of the virus-induced gene-silenced plants revealed a predominant role of *SlPAP26b* over *SlPAP17b* and *SlPAP26a* in the Pi compensation network in tomato seedlings. Transcriptional reporter fusion experiments and transcript profiling studies indicated SlPHR1 and SlPHL1 independent role of *SlPAP26b* in Pi homeostasis in tomato seedlings.

## Materials and Methods

### Plant material, growth conditions, and nutrient deficiency treatments

Tomato seeds (*Solanum lycopersicum* cv. Pusa Ruby, an Indian tomato cultivar) were surface sterilized and incubated in the dark on a petri-plate with half-strength Murashige and Skoog (½ X) MS-agar media for three days in a culture room. On the 3^rd^ or 4^th^ day, the germinated seeds with similar radicle length were transferred to transparent Phyta Jars (polystyrene jars with HDPE lid with overflowing capacity 500 mL, Himedia Pvt. Ltd., India) containing (½ X) MS-agar non-limiting phosphate (HP, 1.25 mM) and grown for next four days. Then, the seedlings were divided into two sets, and one set was transferred again to HP conditions while the other set of seedlings was transferred to limiting/low phosphate (LP, 5 µM) media. The seedlings were grown in a culture room maintained at 25 °C, relative humidity (RH) 60-70%, light intensity ≈200 μmolm^-2 -1^ with 16 h light followed by an 8 h dark photoperiodic regime. LP media was supplemented with an equimolar concentration of KCl to compensate for Potassium. After 8 and 15 days of growth on the respective media, seedlings were harvested and imaged. The length of roots and shoots was measured using ImageJ (https://imagej.nih.gov/ij). Subsequently, root, shoot, and whole seedling tissues were flash-frozen in liquid N_2_ and stored at -80 °C.

Tomato seedlings were grown for 11 days on (½ X) Hoagland’s medium containing (L^-1^): [3 mM KNO_3_, 1 mM MgSO_4_.7H_2_O, 2 mM Ca(NO_3_)._4_H_2_O, 1.25 mM KH_2_PO_4_, 1 mM Fe (III) EDTA, 0.25 mM H_3_BO_3_, 0.002 mM MnSO_4_.H_2_O, 0.002 mM ZnSO_4_.7H_2_O, 0.0005 mM CuSO_4_.5H_2_O, 0.0005 mM Na_2_.MoO_4_ (pH 5.8)] (Hoagland and Arnon, 1950) and then subjected to minerals stress deficiency of Phosphorus (P), Nitrogen (N), Potassium (K), and Iron (Fe) by withdrawing these nutrients, one at a time, form the media. Plants were grown for 15-D post starvation, and the seedlings were stored at –80 °C for RNA isolation and RT-qPCR experiment.

In the second series of experiments related to sucrose (S) deprivation, the seeds were germinated under sterile conditions on (½ X) MS medium supplemented with 1.25 mM phosphate and 2% (58 mM) sucrose. On the fifth day of germination, the sprouted seedlings were transferred to a (½ X) MS medium containing 5 µM phosphate, with or without 2% sucrose. The seedlings were allowed to grow for 8 and 15 days on their respective media: +P+S (1.25 mM phosphate and 2% sucrose), +P–S (1.25 mM phosphate without sucrose), – P+S (5 µM phosphate and 2% sucrose), and –P–S (5 µM phosphate without sucrose).

### Time-course expression kinetics and Pi-recovery experiments

As mentioned in the previous section, the germinated seedlings were transferred on standard (½ X) Hoagland medium. After eleven days, seedlings were subjected to phosphate starvation for the next 15 days in 1L pots containing low phosphate (5 µM) (½ X) Hoagland medium. A set of seedlings were also grown under HP conditions. Shoots and root tissues were harvested from the first day of the starvation in a time-dependent manner (24 h, 48 h, 7D, 15D). During the Pi-recovery experiment, the Pi-starved seedlings were recovered by transferring them from LP to HP media. The samples were collected after 15 min and 4 h of 1.25 mM Pi replenishment. The samples were instantly frozen in liq. N_2_ and stored at -80 °C for further experiments. Further, for remobilization study, node wise leaves were collected from 25-D post starved seedlings for soluble Pi content and molecular study.

### RNA isolation and cDNA synthesis

For RNA-sequencing, total RNA from the pooled tomato seedlings, from three independent experiments performed separately, was isolated using the Total Plant RNA Isolation Kit (Favorgen Biotech, Taiwan). Simultaneously, total RNA from each replicate of pooled seedlings was also isolated separately for the quantitative real-time PCR (RT-qPCR) study. RNase-Free DNase (Promega, USA) treatment of the isolated RNA samples was done to remove any DNA contamination. The quality and quantity of isolated RNA samples were assessed by performing 1% agarose gel electrophoresis and NanoDrop 2000C (Thermo Fisher, USA). The high-quality RNA samples were outsourced for RNA sequencing. One microgram of high-quality RNA was used for the first-strand cDNA synthesis by using an iScript™ cDNA synthesis kit (Bio-Rad Laboratories, USA). Primer specificity was ensured by performing a BLASTN search of each sequence against the *Solanum lycopersicum* ITAGv2.40 database **(Supplementary Table S1).** RT-qPCR reactions were performed in 96-well optical reaction plates (Applied Biosystems, USA) in a PCR cycler CFX96 (Bio-Rad Laboratories, USA). The tomato *GAPDH* gene was used as an internal control in the RT-qPCR analysis (Kumar et al., 2016). For RT-qPCR validation of the transcriptome data, the cDNA synthesized from the same RNA samples sent for RNA sequencing was used as a template. At least two biological and three technical replicates per sample were used in the RT-qPCR analysis of selected candidates. The relative expression level of genes was calculated as described earlier (Kumar et al., 2016).

### RNA-sequencing and data analysis

For RNA-sequencing, the four libraries were sequenced using the Illumina HiSEQ 4000 platform using these parameters: 150 bp paired-end sequencing and ≥ 25 million reads per library. NGSQC Tool kit was used for stringent quality control of the paired-end sequence reads of all the samples. Paired-end sequence reads with a Phred score ≥ Q30 were taken for further analysis. Tomato Genome cDNA (ITAG release v4.0) was used to align the sequenced reads. Kallisto quant and the DESeq R package were used for reads-alignment and differential expression analysis, respectively, with default parameters. Transcriptomes of the same-time point Pi-deficient seedlings were compared with the Pi-sufficient seedlings to establish the differential expression of genes. Transcripts with log_2_ fold change with log_2_ cut-off value of 1 and p-value <=0.05 were considered as significantly differentially expressed genes (DEGs) for respective time points. The unsupervised hierarchical clustering of DEGs was done using Cluster 3.0. The results were visualized using Java Tree View v1.1.6. All the transcript sequences were used for BLAST against the Refseq Plant database to get Gene Ontology (GO) and KEGG pathways for the complete data. Gene ontology (biological processes, cellular components, and molecular functions) for the DEGs was fetched from the Ensemble Plant Biomart (SL3.0; http://plants.ensembl.org/index.html). The raw data is submitted to NCBI-SRA (www.ncbi.nlm.nih.gov/sra) with accession ID SUB8104515.

### Proteome analysis

Tomato seeds were washed and germinated, as mentioned earlier. The germinated seedlings were transferred to (½ X) Hoagland media containing HP (1.25 mM) and LP (5 µM) media. Plants were transferred post-germination to Pi-sufficient and Pi-deficient media and grown for 10 days. The roots of the seedlings were pulverized in Liq. N_2_ and all the samples were lyophilized and subsequently used for proteome analysis by LC/MS. Total protein extracts were prepared as previously described (Jindal et al., 2022) and portions of samples corresponding to 5 µg of peptide were analyzed by nanoflow reverse-phase liquid chromatography-mass spectrometry using a 15 cm C18 Zorbax column (Agilent), a Dionex Ultimate 3000 RSLC nano-UPLC system, and the Orbitrap Fusion Lumos Tribrid Mass Spectrometer equipped with a FAIMS Pro Interface. All samples were analyzed using alternating FAIMS compensation voltages of −40, −50, and −75 V. The measured spectra were recalibrated, filtered (precursor mass—350–5000 Da; S/N threshold—1.5), and searched against the ITAG4.1 (retrieved from the Sol Genomics Network, https://solgenomics.net/ftp//genomes/Solanum_lycopersicum/annotation/ITAG4.1_release/) and common contaminants databases using Proteome Discoverer 2.5 (Thermo, algorithms SEQUEST and MS Amanda (Dorfer et al., 2014). The quantitative differences were determined by Minora, employing precursor ion quantification followed by normalization (total area) and calculation of relative peptide/protein abundances. The mass spectrometry proteomics data have been deposited to the ProteomeXchange Consortium via the PRIDE (Perez-Riverol et al., 2022) partner repository with the dataset identifier PXD044390.

### Non-targeted metabolite profiling

For gas chromatography-mass spectrometry (GC-MS) based profiling of metabolites, 8-D and 15-D HP- and LP-grown tomato seedlings in half-strength (½ X) MS-agar media, were frozen and powdered in liq. N_2_. 150 mg of each frozen sample was used for extraction in 1400 μL of 100 % MeOH solution. The further processing of the extracted samples was performed by strictly following the steps described previously by (Sharma et al., 2020). Ribitol (0.2 mg ribitol/mL water) acted as an internal standard. Gas chromatography was performed using a hybrid quadrupole time-of-flight instrument (Leco Corporation, USA). Processing and initial analysis of the raw data was done using the ChromaTOF and MetAlign packages (Lommen and Kools, 2012). The mass signals present in at least three samples were used for further research. The metabolites were identified using the NIST (National Institute of Standards and Technology) GC/MS Metabolomics Library software (NIST/EPA/NIH Mass Spectral Library 14, v 2.2, Department of Commerce, USA). PCA and statistical analysis were performed in MetaboAnalyst 4.0 (Xia et al., 2012). At least five biological replicates per sample were used in the metabolic profiling.

### Quantification of total soluble Pi content and total P

To quantify the total soluble Pi, 60 mg of fresh tissue was powdered in a prechilled pestle and mortar. The powder was mixed with 250 µL of 1 % glacial acetic acid in 1.5 mL microcentrifuge tubes (MCTs), and the solution mixtures were thoroughly mixed by vortexing. Samples were incubated in liquid N_2_ for 30 sec and then left at room temperature for thawing. The thawed samples were centrifuged at 5000x*g* for 10 min at room temperature (RT). 50 μL supernatant from MCT was subjected to soluble Pi quantification using a phosphomolybdate colorimetric assay, as described by (Ames, 1966). Each experiment was independently repeated at least three times, with 20 or more tomato seedlings per repeat.

For total P estimation, whole seedlings were dried at 65 °C using a hot air oven for 24 h. The dried seedlings were ground to a fine tissue powder, and 50 mg of powdered tissue was taken in a 100 mL boiling tube. 1 mL of concentrated sulfuric acid was added in drops to each tube, and the mixture was incubated overnight at RT. The tubes were transferred to a hot air-oven the next day, maintained at 120 °C for 2 h. Subsequently, 30 % H_2_O_2_ was added dropwise to each hot tube followed by incubation at RT for 30 min until the mixture became colorless. The 10 μL solution from each tube was transferred to a new 1.5 mL MCT (600 μL of 0.42 % acidic ammonium molybdate and 100 μL of 10 % ascorbic acid). After mixing properly, the MCT was kept at 45 °C for 20 min. Later, 200 μL solution was taken from each tube for the absorbance at O.D. 820 nm. Total P was quantified using a P standard curve (Srivastava et al., 2021b). Each experiment was independently repeated at least three times, with ten or more tomato seedlings per repeat.

### Root-associated secretory acid phosphatase (SAP) and intracellular acid phosphatase (IAP) activity assays

For root-associated SAP activity, roots from Pi-sufficient and Pi-deficient tomato seedlings were excised and washed carefully with double distilled water. The roots were placed in a para-Nitrophenylphosphate (pNPP) containing reaction buffer (10 mM MgCl_2_, 50 mM NaOAc, pH 4.9). After incubation at 37 °C for 4 h, the reaction was terminated by adding 100 μL of 0.4 N NaOH solution. The roots were then removed from the setup, and the absorbance of the remaining solution was recorded at 410 nm. Root-associated total secretory APase activity is expressed as A_410nm_/ root/h (Srivastava et al., 2020).

For IAP, total protein was extracted from root and leaf tissues in an ice-cold extraction buffer, as described by (Lu et al., 2016). The protein concentration was quantified using a Bradford reagent. 10 µg of total protein was incubated with 590 µL reaction buffer containing (10 mM MgCl_2_, 50 mM NaOAc, pH 4.9). Subsequently, 80 μL of pNPP was added to the reaction buffer. Reactions were incubated at 37 °C for 1 h and stopped by adding 120 μL 0.4N NaOH. The absorbance was determined at 410 nm using a microplate assay (TECAN, https://www.tecan.com). The APase activity was expressed as A_410nm_/mg protein/h (Wang et al., 2011). The experiment was independently repeated thrice.

Total APase isoforms in root and shoot samples were detected separately using In-gel assay. In brief, shoot and root tissues were grounded to extract the crude protein using protein extraction buffer (0.1 M KOAc, 20 mM CaCl_2_, 0.1 mM PMSF, pH 5.5). Protein concentration was calculated using Bradford assay reagent. The isozyme profile of APases was visualized by loading 10 µg crude protein on SDS-PAGE (denaturing gel). Gel was stained using chromogenic substrates solution conatining, β-NAP and Fast black K, as described by Wang et al. (2014) (Wang et al., 2014).

### Quantification of total carbohydrates

The total carbohydrates were quantified using the phenol-sulfuric acid method (HiPer Carbohydrates Estimation Teaching Kit-Himedia). First, we plotted the glucose standard curve with the standard glucose (1 mg/mL). Fresh tissue was ground in liq. N_2_ and 10 mg powdered tissue were transferred in 2 mL MCTs. Afterward, 0.2 mL of 0.5 % phenol solution was added to all the samples, followed by gentle mixing. Then, 1 mL of concentrated sulphuric acid was added to each MCT. Samples were mixed well, and incubated for 10 min at RT, followed by a second incubation at 30 °C for 20 min. Finally, the O.D. was recorded at a wavelength of 490 nm. Total soluble sugars were estimated by using the standard plot made by using glucose (1 mg/ml). For the estimation of sucrose, fructose, and glucose, a Megazyme kit (Catalog No # K-SUFRG, Lot 210733-6) was used. The samples were processed as per the manufacturer’s protocol.

### Estimation of total anthocyanins

Total anthocyanin pigments were quantified from leaf tissue. For that, 150 mg of shoot tissue was grounded in 10 mL of acidic methanol (methanol: water: HCl, 79:20:1, v/v/v), and the extraction was kept in the dark for 2 h. After incubation, samples were centrifuged at 5000 x*g* at RT for 10 min. The absorbance of the supernatant was measured at A_530_ nm and A_657_ nm. Total anthocyanins content was measured using the following formula: {A_530_-(A_657 nm_/3)}/gm FW where A_530 nm_ and A_657 nm_ are absorbances in nm of the extracted pigments solution (Jiang et al., 2007). The experiment was independently performed three times.

### Virus-induced gene silencing (VIGS)

Gene fragments of *SlPAP17b, SlPAP26a, SlPAP26b,* and *SlPDS* for virus-induced gene silencing experiments were cloned in the *pTRV2* vector and infiltrated in the tomato seedlings, as described earlier (Akash et al., 2022; Singh et al., 2023). *Agrobacterium tumefaciens GV3101* strain was used in the VIGS experiments. Fully opened cotyledons of seven-day-old tomato seedlings of the Pusa Ruby cultivar were used for agroinfiltration. *pTRV1/pTRV2* plasmids infiltrated plants were used as empty vector (EV) controls.

### Western blot analysis

500 mg of fresh tissue collected from tomato seedlings was powdered in a prechilled pestle and mortar. The powder was suspended in prechilled protein extraction buffer (100 mM Tris-base pH 7.5, 10% Glycerol, 100 mM EDTA-Na_2_, 2 mM EGTA, 2 mM PMSF, and 1X-protease arrest) and incubated on ice for 30 min. The samples were centrifuged at 24000x*g* for 30 min at 4 °C, and the supernatant was collected. The crude protein was quantified using Bradford’s method, as described previously. 20 μg protein was used for western blot analysis using SlSPX2 polyclonal antibody conjugated with KLH (Keyhole Limpet Hemocyanin), as described previously (Singh et al., 2023).

### *SlPAP26bpro::GUS* transcriptional activation assay and histochemical staining

The 1.5-kb upstream promoter region of *SlPAP26b* was PCR amplified. The amplified product was cloned in the *pCAMBIA1391z* binary vector. The cloning was confirmed by Sanger sequencing. The confirmed plasmids were mobilized into the *Agrobacterium tumefaciens EHA101* strain. The recombinant agrobacterium cultures were used to infiltrate the top leaves of 6–8-week-old tobacco plants. In brief, confirmed single colonies from each vector were inoculated into liquid Yeast Mannitol media (YEM) containing kanamycin (50 mg/L) and rifampicin (30 mg/L) at 28 °C for 24 h. The secondary culture was initiated from primary cultures in 10 mL YEM supplemented with kanamycin (50 mg/L), rifampicin (30 mg/L), and 200 μM acetosyringone for 24 h at 28 °C. The bacterial cells were pelleted by centrifugation at 3000x *g,* and pellets were resuspended in MMA (10 mM MES pH 5.6, 10 mM MgCl_2_, 200 μM acetosyringone) with a final OD_600 =_ 1. The resuspended cultures were incubated for 2 h at room temperature. The cultures containing different plasmids were mixed in 1:1:1 (*pCAMBIA1391z-SlPAP26bpro:GUS::pCAMBIA1302:SlPHR1/SlPHL1::p19*) or (*pCAMBIA1391z-GUS::pCAMBIA1302*:*SlPHR1/SlPHL1*::*p19*) ratio. Control plants were infiltrated with blank vectors. The infiltrated plants were first kept in the dark for 24 h and later shifted to a culture room maintained at 22 °C with RH-65 %, light intensity 200 μmol/m^2^/s^-1^ with 16/8 h photoperiodic cycle for 72 h. For histochemical analysis, the infiltrated leaves were removed from plants and gently washed. These leaves were submerged in the GUS solution [(50 mM NaH_2_PO_4-_Na_2_HPO_4_, pH 7.3, 2 mM K_3_Fe (CN)_6_, 2 mM K_4_Fe (CN)_6_, 1 mM 5-bromo-4-chloro-3-indolyl-β-D-glucuronide (X-Gluc), 0.1 % (v/v) Triton X-100)] at 37 °C for overnight. The leaves were then treated with a clearing solution (95% ethanol) to remove chlorophyll (Gambhir et al., 2022). The cleared leaves were then photographed.

### Statistical analysis

Each experiment was performed using a minimum of three biological replicates and presented as mean values after calculating the standard deviation. Error bars denote standard deviation. A statistically significant difference between the control and treated samples at a 5% level was evaluated by Unpaired T-test and Two-Way ANOVA [(p-value < 0.12 (ns), 0.033 (*), 0.002 (**), 0.001(***)]. Significance as indicated by an asterisk.

## Results

### Activation of multiple acid phosphatases contributes to the enhanced acid phosphatase activity in Pi-deficient tomato seedlings

Several PAPs have been characterized for different roles in plant (Bhadouria and Giri 2022). For example, *AtPAP15*, a purple acid phosphatase with phytase activity, has been implicated in a lower in-vitro germination rate of the pollen in Arabidopsis (Kuang et al., 2009). The root-associated *AtPAP10* and its homologs are involved in plant tolerance to Pi limitation (Deng et al., 2020; Suen et al., 2015; Wang et al., 2011). AtPAP26 homologs play a pivotal role in extracellular phosphate-scavenging or Pi remobilization (Robinson et al., 2012; Gao et al. 2017). Because P starvation responses, such as in root system architecture and acid phosphatase activity, may vary in different genetic backgrounds of tomato (Demirer et al., 2023; Satheesh et al., 2022; Srivastava et al., 2020), we first studied the response of Pusa Ruby seedlings under low Pi availability at two-time points, i.e. 8-days (8-D) and 15-days (15-D) **(Fig. 1A-E).** As expected, Pi-starvation led to a severe reduction in the total soluble Pi content at both 8-D and 15-D in the Pi-deficient seedlings **(Fig. 1B).** Longer primary roots, altered number of lateral roots number, enhanced total carbohydrate levels, and higher total anthocyanins accumulation were also observed in Pi-deficient seedlings than their controls at 8-D and 15-D time points **(Fig. 1E, Supplementary Fig. Sf1).** Contrary to a moderate but significant increase in the total IAP activity (which increased by 13%) at 15-D, a more pronounced increase in the SAP activity at both time-points (which increased by 23% at 8-D and by 46% at 15-D) was noticed in Pi-deficient seedlings **(Fig. 1C, D)**.

**Fig. 1:**
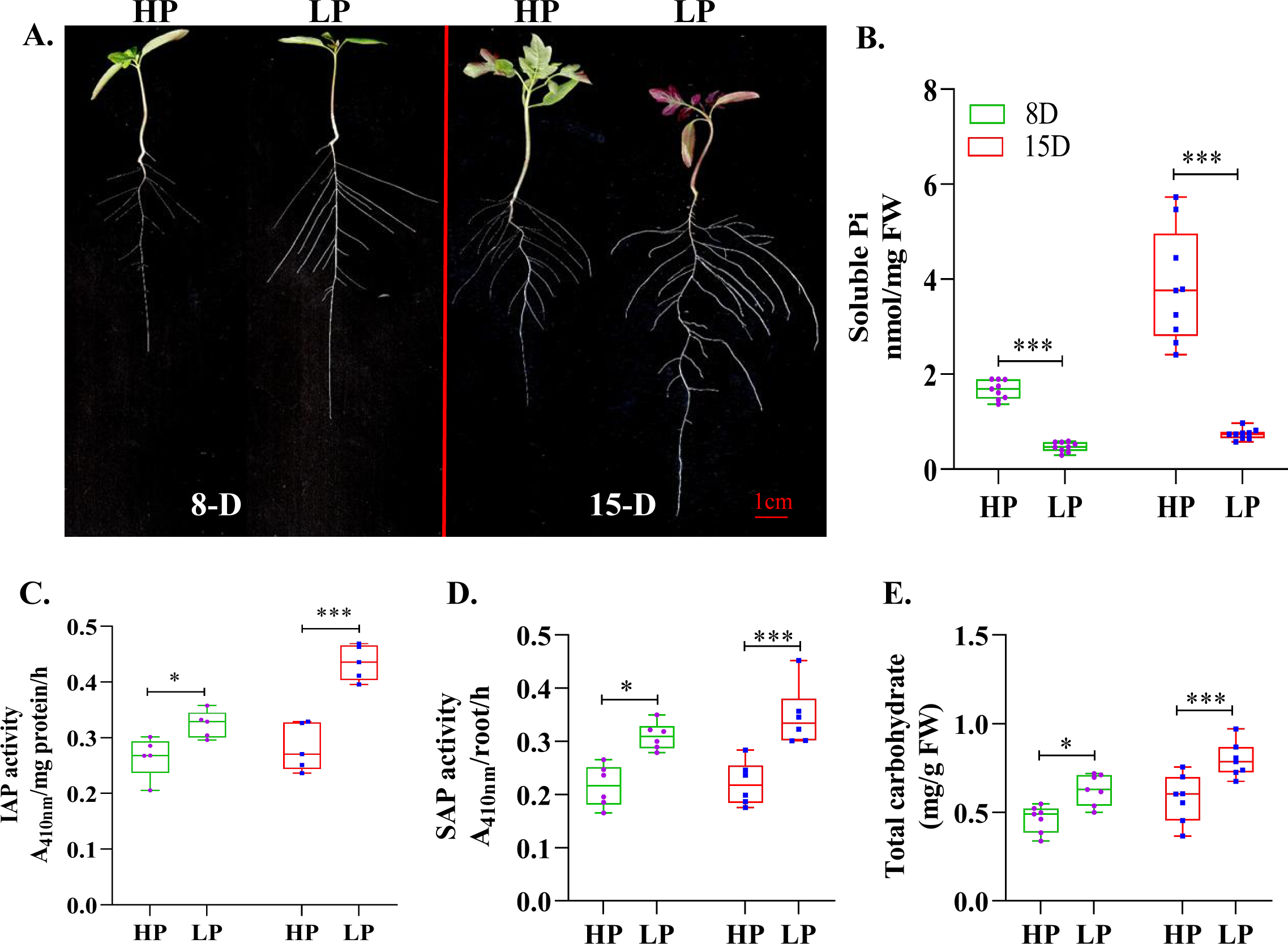
Physio-biochemical characterization of tomato seedlings grown in (½ X) MS agar media containing HP (1.25 mM; Pi-sufficient) and LP (5 µM; Pi-deficient) concentration for 8-days (8-D) and 15-days (15-D) after germination. **(A)** Images of 15-D old tomato seedlings. **(B)** Estimation of total inorganic phosphate (Pi) content. **(C-D)** Intracellular acid (IAP) and secretory acid phosphatase (SAP) activity. **(E)** Total carbohydrate content. The experiment was performed in three independent biological replicates. (n=30). HP, high phosphate; LP, low phosphate. The scale bar denotes 1 cm. Asterisks represent the statically significant differences based on two-way ANOVA tests [(p-value < 0.12 (ns), 0.033 (*), 0.002 (**), 0.001 (***)]. Error bars indicate standard deviation.

We next studied transcriptomes of Pi-sufficient and Pi-deficient tomato seedlings to identify the complete set of acid phosphatases contributing to the observed enhanced APase activity in our biochemical assays. The differential analysis of RNA-sequencing data (four libraries ∼30 million high-quality reads with >= 30 Phred score with an excellent alignment percentage of over 90 % to the reference tomato genome; **Supplementary Table S2)** identified 1246 and 1201 transcripts as differentially expressed genes (DEGs) at 8-D and 15-D time points, respectively **(Supplementary Table S3)**. In comparison to the 669 upregulated transcripts at 8-D, mRNA levels of 922 genes were elevated at 15-D **(Fig. 2A)**. The analysis of common DEGs, shared by Pi-deficient seedlings at both the time-points, revealed upregulation of 316 common transcripts **(Fig. 2A)**. Among these, 15 transcripts corresponded to acid phosphatases, including several purple acid phosphatases such as *SlPAP1* (Solyc05g012260), *SlPAP10b* (Solyc01g110060), *SlPAP15* (Solyc09g091910), *SlPAP17b* (Solyc03g098010), *SlPAP26a* (Solyc12g009800) and *SlPAP26b* (Solyc07g007670) **(Fig. 2C)**. Other acid phosphatases, corresponding to Solyc01g095980, Solyc04g008250, Solyc06g062390, and Solyc06g062380 loci were also induced in the Pi-deficient seedlings. The accuracy of RNA-sequencing data was validated by real time quantitative PCR (RT-qPCR) analysis of 17 genes. The high correlation observed in the expression profiles of 14 genes between RT-qPCR and RNA-sequencing data at both time points supported the differential gene analysis results observed in the transcriptome analysis **(Supplementary Fig. Sf2A, B)**. Further, many of these acid phosphatases, with elevated transcript levels, were also upregulated at the proteome level. Higher to moderate peptide abundance of *SlPAP1*, *SlPAP10a* (Solyc01g110050), *SlPAP10b, SlPAP15, SlPAP17b, SlPAP26b*, and *SlPAP27b* (Solyc07g007670) confirmed the contribution of multiple acid phosphatases to the observed enhanced APase activity in Pi deficient seedlings **(Fig. 2D).**

**Fig. 2:**
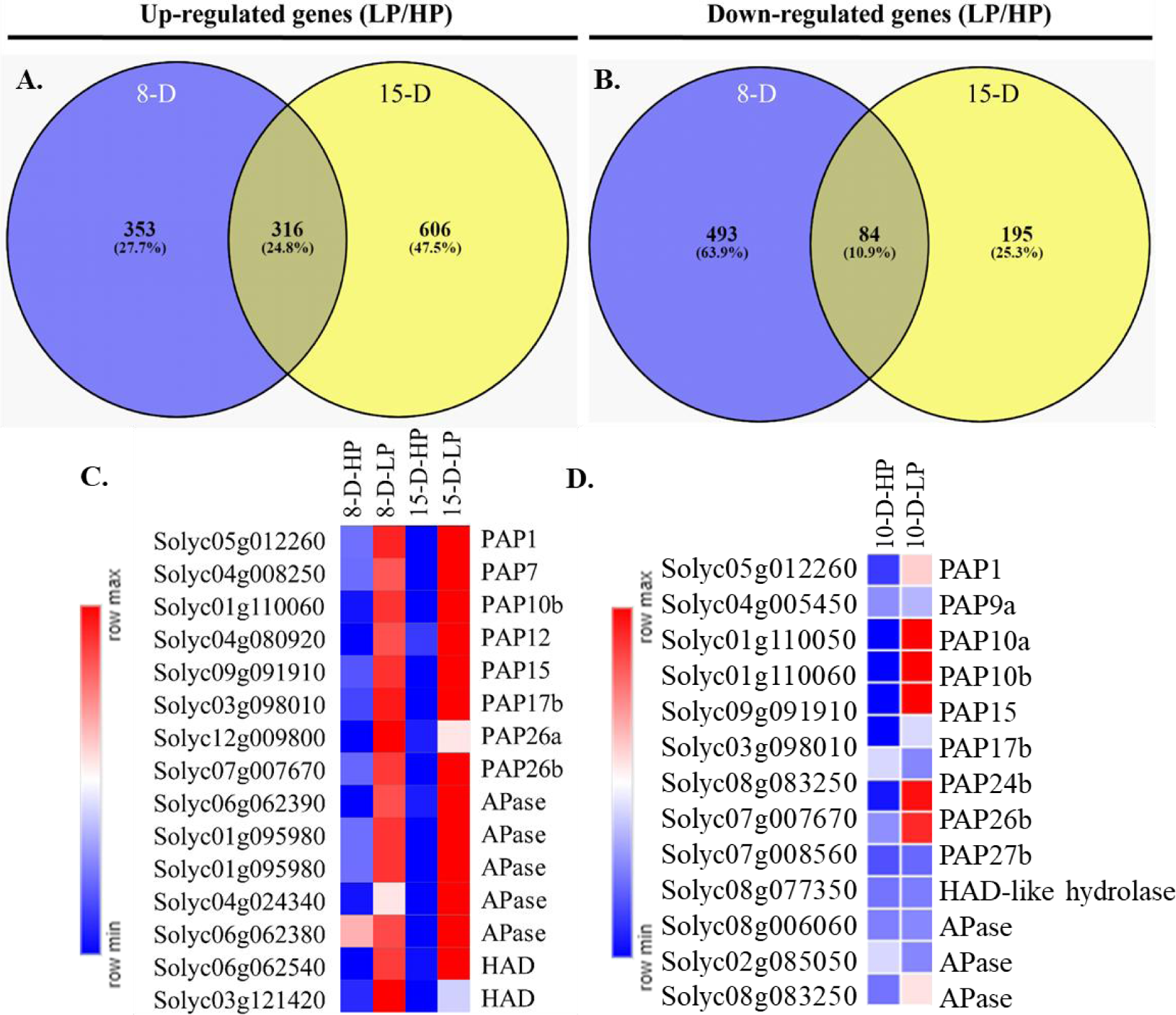
Overview of global transcriptional changes in Pi-starved seedlings. **(A, B)** Venn diagram representing the common as well as unique differentially expressed genes (DEGs) identified in the RNA-sequencing data at 8-D and 15-D of Pi starvation. **(C-D)** Transcript and protein abundance of acid phosphatase genes in transcriptome **(C)** and proteome **(D)** data, respectively. Heatmaps were generated using mean base values obtained for each gene using Morpheus (https://software.broadinstitute.org/morpheus/) with hierarchical clustering, euclidean metric, and average linkage method. Error bars indicate standard deviation. HP; Pi sufficient, LP; Pi-deficient.

### External sucrose supply influences acid phosphatase activity in tomato seedlings

Pi deprivation reduces shoot growth and enhances root proliferation, activates PSI genes such as purple acid phosphatases, phosphodiesterases, and sugar transporters, and invigorates P starvation responses such as increased APase activity by allocating photosynthates to roots in Pi-deficient plants (Hammond and White, 2008; Karthikeyan et al., 2007; Müller et al., 2007). Mutation in *SUCROSE2* gene, phloem-specific sucrose (Suc)/H+ symporter is involved in phloem loading and efficient sucrose transport from source tissues to sink tissues in Arabidopsis, inhibits APase activity by 30% in the mutant seedlings than wild-type plants (Zakhleniuk et al., 2001). Because enhanced total carbohydrates levels were observed in the Pi-deficient seedlings, we next performed untargeted metabolites analysis of Pi-deficient and Pi-sufficient seedlings at both time points **(Fig. 3).** This analysis clearly showed enrichment of several sugar metabolites such as fructopyranose, D-psicofuranose, sucrose and glucopyranose, in the Pi-deficient seedlings compared with Pi-sufficient counterparts at 15-D time point and confirmed the elevated levels of carbohydrates in tomato seedlings under Pi deficiency **(Fig. 3; Supplementary Fig. Sf3).** Besides sugar-metabolites, organic acids such as fumaric acid, malic acid, and α-ketoglutaric acid showed a more prominent enhancement in their accumulation at the 15-D time point, indicating a more severe PSR at this time point in the Pi-deficient tomato seedlings **(Fig. 3).**

**Fig. 3:**
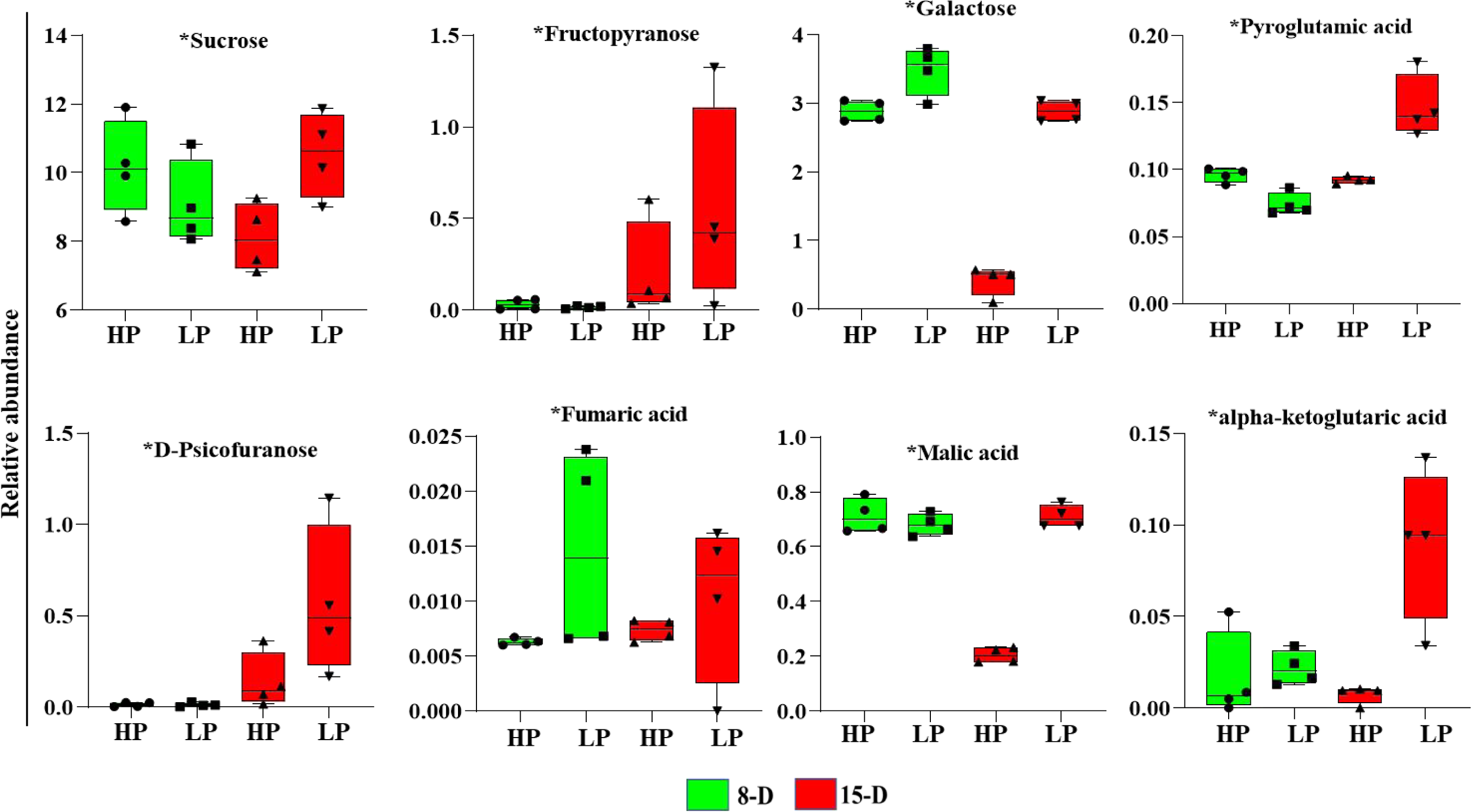
Metabolite profiling of Pi-starved tomato seedlings at 8-D and 15-D of Pi starvation. Box plots representation of selected metabolites. Error bars indicate standard deviation. HP; Pi-sufficient, LP; Pi-deficient. * = denotes significant abundance of detected metabolites.

We next examined the effect of exogenous sucrose supplementation on the P starvation response in tomato seedlings. For this, five-day-old seedlings (1.25 mM phosphate and 2 % sucrose) were transferred onto Pi-sufficient/Pi deficient (½ X) MS medium with or without sucrose (2 %) and grown for 8 and 15 days **(Fig. 4; Supplementary Fig. Sf4).** At both time points, the soluble Pi content was significantly decreased in –P+S seedlings as compared to +P+S, +P–S, and –P–S seedlings **(Fig. 4A).** Sucrose supply increased SAP and total root surface associated acid phosphatase activities in +P+S and –P+S seedlings at both time points compared to +P–S and –P–S seedlings **(Fig. 4B, C).** The elevated mRNA levels of *LePS2,* a previously characterized PSI gene with acid phosphatase activity (Baldwin et al., 2008), and selected PSI PAPs identified in this study, such as *SlPAP17b* and *SlPAP26b,* in – P+S seedlings than –P–S or +P–S counterparts supported the positive role of sucrose on the activation of these genes and enhanced acid phosphatase activity in Pi-deficient tomato seedlings at the molecular level. Interestingly, *SlPAP26b* transcriptional activation was found to be independent of the plant’s Pi status, as transcripts of this gene increased significantly in the presence of sucrose even under Pi-sufficient conditions over their respective controls **(Fig. 4D)**. Further, the unaltered transcript levels of *SlPAP26a* in this experiment indicate that not all PSI genes respond positively to sucrose. *SlPT1* and *SlPT7* genes were used as positive controls in the RT-qPCR analysis **(Fig. 4D)**.

**Fig. 4:**
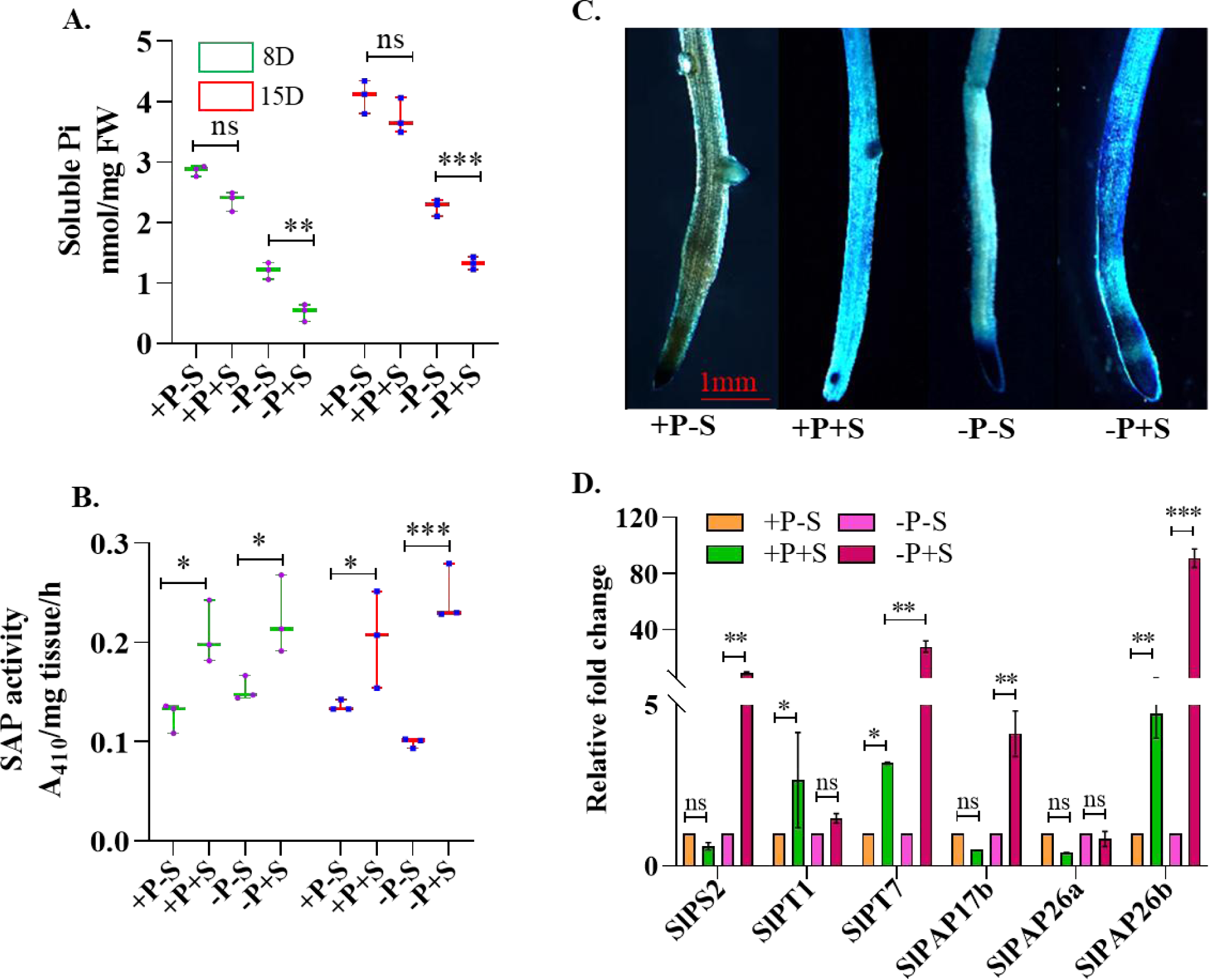
Effect of sucrose on P starvation response in tomato seedlings under HP (Pi-sufficient) and LP (Pi-deficient) conditions. **(A)** Total soluble Pi content level in with and without sucrose tissues during HP and LP conditions. **(B, C)** Secretory acid phosphatase activity (SAP) by pNPP assay and BCIP staining for root-associated APase activity. **(D)** Expression analysis of some, selected PSI genes upon sucrose treatment. +P-S = (HP without sucrose), +P+S (HP with sucrose), -P-S (LP without sucrose), -P+S = (LP with sucrose). Asterisks represent the statistically significant differences based on two-way ANOVA [(p-value < 0.12 (ns), 0.033 (*), 0.002 (**), 0.001 (***)]. Error bars indicate standard deviation.

### *SlPAP17b* and *SlPAP26b* are prominently activated in older senescing leaves

Due to the higher demand of Pi in shoot during different biosynthetic processes, we next investigated the expression profiles of selected SlPAPs during Pi remobilization (**Supplementary Fig. Sf5**; Srivastava et al., 2020). For this purpose, we first estimated the total soluble Pi levels in leaves from different nodes, including node 1 (the oldest leaf showing symptoms of senescence), node 3 (middle green leaf), and node 6 (The youngest green leaf) of Pi-deficient and Pi-sufficient tomato plants. This analysis clearly showed the lowest Pi levels in the oldest senescing leaves, followed by node 3 leaf **(Fig. 5A and B).** The youngest leaf (node 6) of both Pi-sufficient and Pi-deficient plants accumulated the highest Pi levels among the leaves used in the study **(Fig. 5B).** This incremental increase in Pi levels from the oldest to the youngest leaf indicates active remobilization of Pi in these tomato seedlings. The following RT-qPCR analysis of SlPAPs, the commonly upregulated members identified in the transcriptome and proteome data with high transcript levels in the aerial parts of the tomato plants, *SlPAP1, SlPAP10b, SlPAP17b, SlPAP26b*, and *SlPAP26b* revealed higher expression of *SlPAP17b* and *SlPAP26b* in these leaves under both Pi-sufficient and Pi-deficient conditions, implicating these two members at the forefront of Pi remobilization in tomato seedlings **(Fig. 5C).** A vacuolar PHT5 class transporter, *SlPHT5;1* was used as a positive control in the RT-qPCR analysis **(Fig. 5D).** In this analysis, *SlPAP26b* and *SlPHT5*;*1* showed similar expression pattern, gradually decreased from older to younger leaves, in Pi-sufficient and Pi-deficient plants **(Fig. 5C, D)**.

**Fig. 5:**
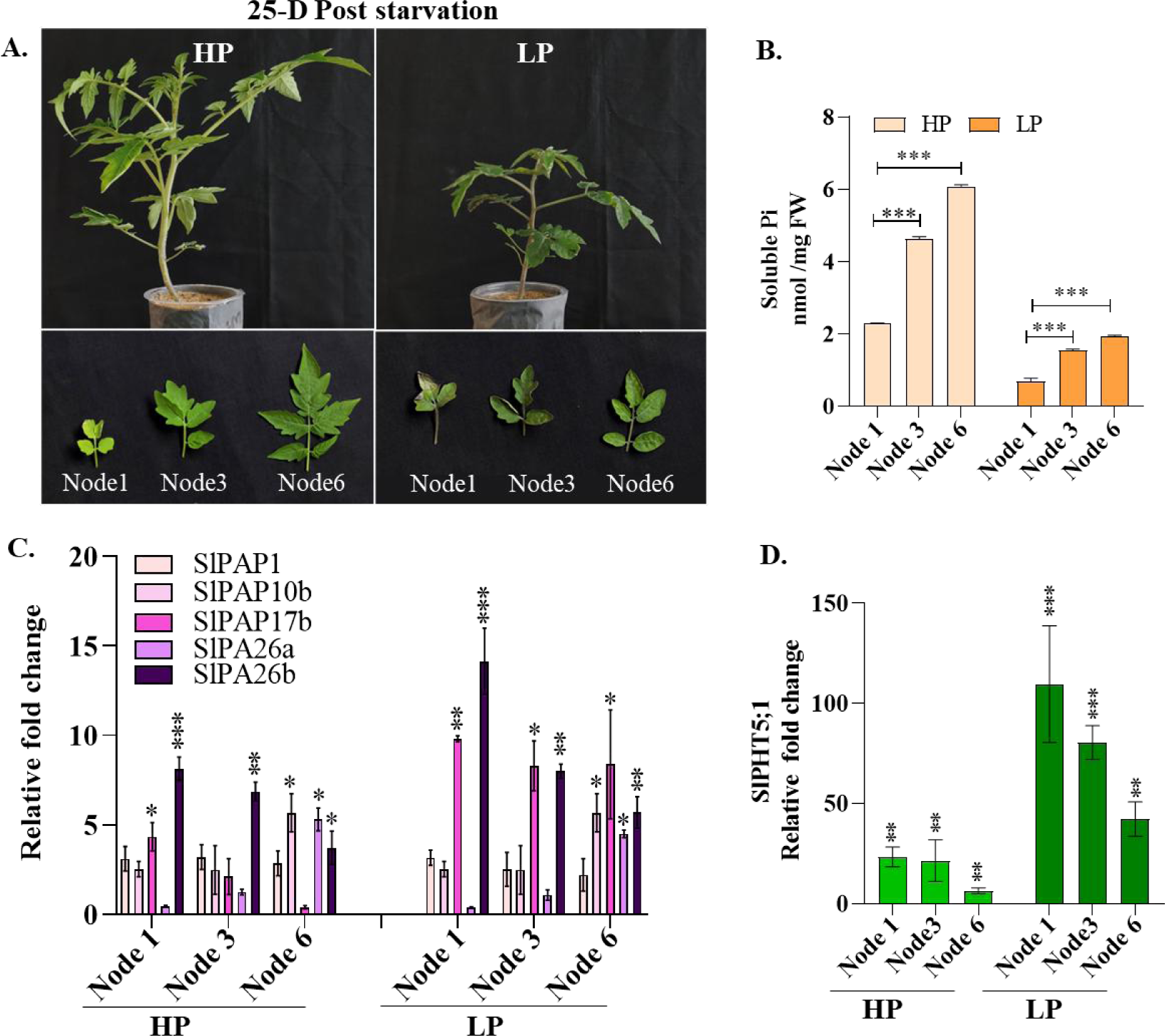
Phosphate remobilization and transcript level of selected purple acid phosphatase candidates in tomato seedlings. **(A, B)** Tomato seedlings image and soluble Pi contents in the leaves from nodes 1, 3, and 6. The analysis was done 25 days post-Pi starvation. **(C)** Expression pattern of PSI PAPs, including *SlPAP1*, *SlPAP10b*, *SlPAP17b*, *SlPAP26a*, and *SlPAP26b* in the leaves of different nodes. These genes have high transcript levels in the shoot and were identified as PSI genes in our transcriptome and proteome data. **(D)** Transcript profiling of *SlPHT5;1*, a close homolog of vacuolar Pi efflux transporter *OsPHT5;1* gene, in node1, node3, and node6 leaves of 25-days Pi starved or same age control HP seedlings. *SlPHT5;1* served as a positive control for SlPAPs used in this experiment. Asterisks represent the statistically significant differences based on two-way ANOVA tests [(p-value < 0.12 (ns), 0.033 (*), 0.002 (**), 0.001 (***)]. Error bars indicate standard deviation.

### *SlPAP17b* shows stronger and more rapid induction upon Pi starvation than *SlPAP26* isoforms

Among the well-characterized Arabidopsis PAPs, evidence shows the predominant role of *AtPAP17* and *AtPAP26* in maintaining Pi-homeostasis by regulating the utilization and mobilization of intracellular or extracellular Pi (Farhadi et al., 2020; Robinson et al., 2012; Wang et al., 2014). In this study, the transcriptome and proteome data confirmed the PSI nature of *SlPAP17* (only *SlPAP17b*) and *SlPAP26* isoforms. The stronger induction of these genes in the older senescing leaves further indicated plausibly similar roles played by these genes in Pi nutrition in tomato seedlings. However, the functions of these genes in Pi homeostasis remained uncertain, and hence, their isoforms were selected for further characterization. Analysis of publicly available RNA-sequencing data (Sato et al., 2012) revealed similar expression profiles for *SlPAP17a* and *SlPAP17b* isoforms, highly restrictive to floral tissue-specific, with almost undetected transcript levels in the leaves (**Supplementary Fig. Sf5).** Due to the non-PSI nature of *SlPAP17a* isoform and its almost identical expression profile to that of *SlPAP17b*, *SlPAP17a* was not included in further experiments. In contrast, *SlPAP26* isoforms (*SlPAP26a* and *SlPAP26b*) exhibited distinct transcript abundance profiles during tomato development. While *SlPAP26b* is strongly expressed in all organs, its paralog *SlPAP26a,* was found to be expressed weakly throughout plant growth and development **(Supplementary Fig. Sf5).** The RT-qPCR analysis of *SlPAP17b*, *SlPAP26a,* and *SlPAP26b* in the Indian cultivar further corroborated with their RNA-sequencing based transcripts profiles **(Fig. 6A)**. We next tested the specificity of the PSI nature of these selected PAPs by examining their transcript levels in the tomato seedlings subjected to deficiencies of other nutrients, including nitrogen (N), potassium (K) and iron (Fe) for 15 days. Unaltered transcript levels of these genes in the deficiency of other mineral nutrients (–N, –K, or –Fe) supported the PSI nature of *SlPAP17b* and *SlPAP26b* isoforms **(Fig. 6B).**

**Fig. 6:**
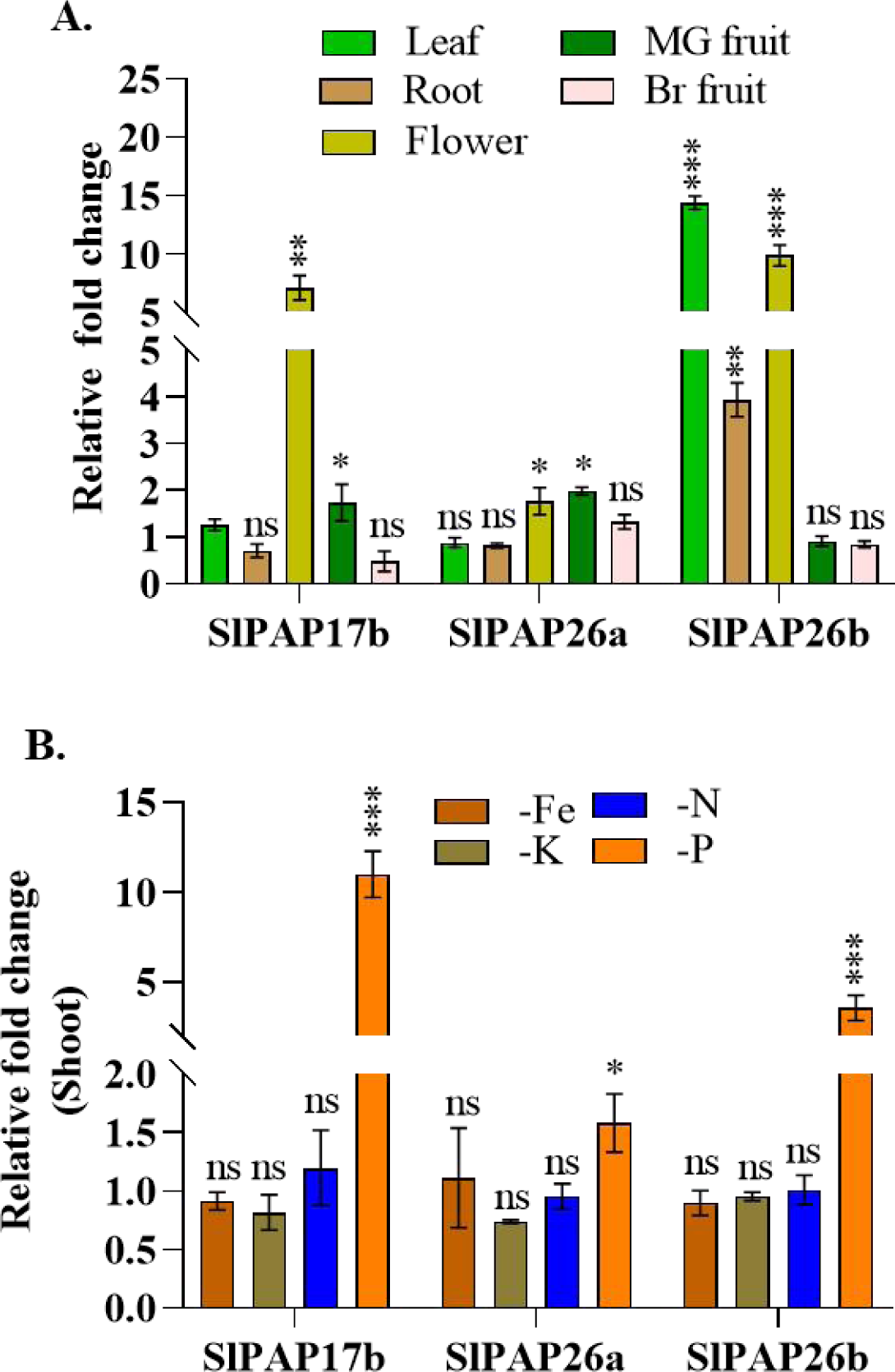
RT-qPCR profiling of *SlPAP17b*, *SlPAP26a,* and *SlPAP26b* during tomato development and under the deficiency of –Fe, –N, –K nutrients. **(A)** Different tissues/stages. The relative expression of these genes in the tissues/stages was determined against their expression in two-week-old hydroponically grown tomato seedlings. **(B)** under the deficiency of mineral nutrients such as iron (Fe), nitrogen (N), or potassium (K). Pi deficiency was used as a positive control in this experiment. Tomato seedlings were subjected to the deficiencies of N, K, or Fe for 15 days in the same way as done for Pi starvation. The relative fold change expression of selected PAPs in the nutrients deficient plants was calculated against their mRNA abundance in Pi-sufficient controls. Asterisks of fig. A & B represent the statistically significant differences based on two-way ANOVA tests [(p-value < 0.12 (ns), 0.033 (*), 0.002 (**), 0.001 (***)]. Error bars indicate standard deviation.

The time course kinetics of the transcript accumulation of these three genes revealed more rapid induction of *SlPAP17b* in both shoot and root tissues of Pi-deficient plants. While *SlPAP26a* mRNA levels increased moderately in the shoot, *SlPAP26b* transcripts were induced in the root at 15-D and in the shoot of Pi-deficient seedlings at 8-D and 15-D time points **(Fig. 7A, B)**. The stronger induction of *SlPAP26a* and *SlPAP26b* transcripts in shoots than roots implies the predominant role of these acid phosphatases in the aerial part of the plants **(Fig. 7A, B).** In the Pi resupply experiments, the transcript levels of all three genes recovered to normal levels within 4 h of Pi resupply, confirming their PSI nature **(Fig. 7A, B).**

**Fig. 7:**
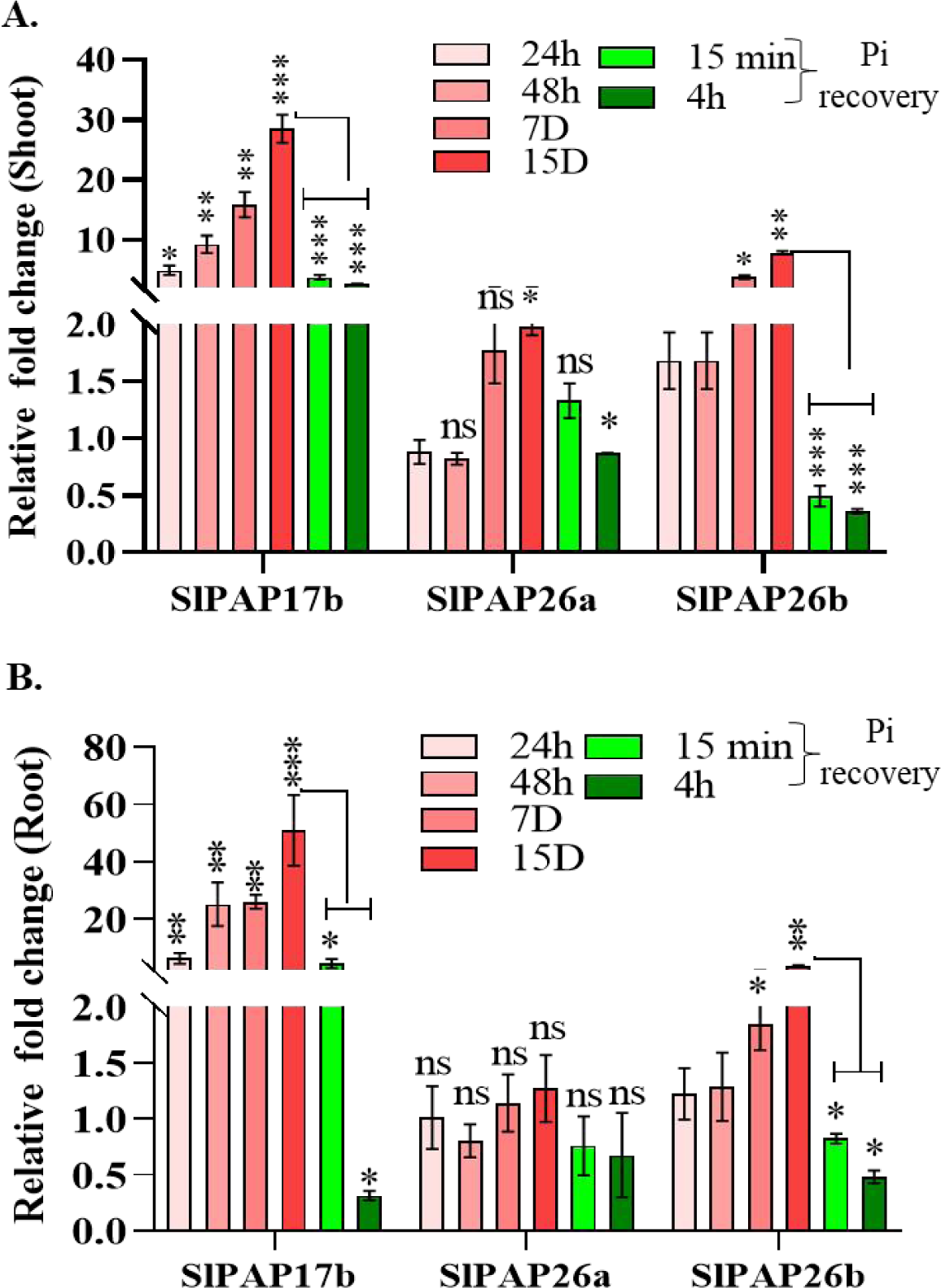
Time-course study of the transcript levels of *SlPAP17b*, *SlPAP26a,* and *SlPAP26b* genes during Pi-starvation and Pi recovery. Time course gene activation kinetics and recovery of these genes in Pi-starved shoot **(A)** and root tissues **(B).** The relative fold expression of these genes at different Pi-starvation time points was calculated against their transcript levels in the unstressed counterparts at the start of Pi withdrawal. For 15 min and 4 h Pi recovery samples, 15-D Pi starved seedlings were used as the control to calculate the relative fold change. Asterisks represent the statistically significant differences based on two-way ANOVA tests [(p-value < 0.12 (ns), 0.033 (*), 0.002 (**), 0.001 (***)]. Error bars indicate standard deviation.

### *SlPAP26b* encodes a major purple acid phosphatase as its silencing exacerbates the P starvation response in tomato seedlings

To investigate the role of *SlPAP17b*, *SlPAP26a,* and *SlPAP26b* in Pi starvation, we next transiently silenced these genes using Virus-induced gene silencing (VIGS). In VIGS experiments, *SlPDS* silencing was used as a positive control. The photobleached leaf phenotype observed in most of the infiltrated plants (≥80% across the experiments) confirmed the highly effective gene-silencing in VIGS experiments **(Supplementary Fig. Sf6 A, B, C).** At molecular levels, RT-qPCR analysis confirmed the inhibited transcript levels of candidate SlPAPs, by at least 60%, in *SlPAP17b, SlPAP26a,* and *SlPAP26b* infiltrated tomato seedlings compared to their EV infiltrated controls **(Fig. 8A, B, Supplementary Figs. Sf7 A, B and Sf8 A, B).** *SlPAP17b* and *SlPAP26a* silenced seedlings failed to exhibit any phenotypic difference vis-à-vis their EV controls. The biochemical assays for total P and total anthocyanins also did not highlight significant differences between *SlPAP17a* and *SlPAP26a* silenced seedlings and the EV controls **(Supplementary Figs. Sf7, Sf8).** However, total soluble Pi content decreased by 29% in *SlPAP17b* silenced seedlings under Pi-sufficient conditions. These seedlings also exhibited reduced phosphorus use efficiency (PUE), which decreased significantly by 13% and 29% under both Pi-sufficient and Pi-deficient conditions to their EV controls, respectively **(Supplementary Fig. Sf8, F).**

**Fig. 8:**
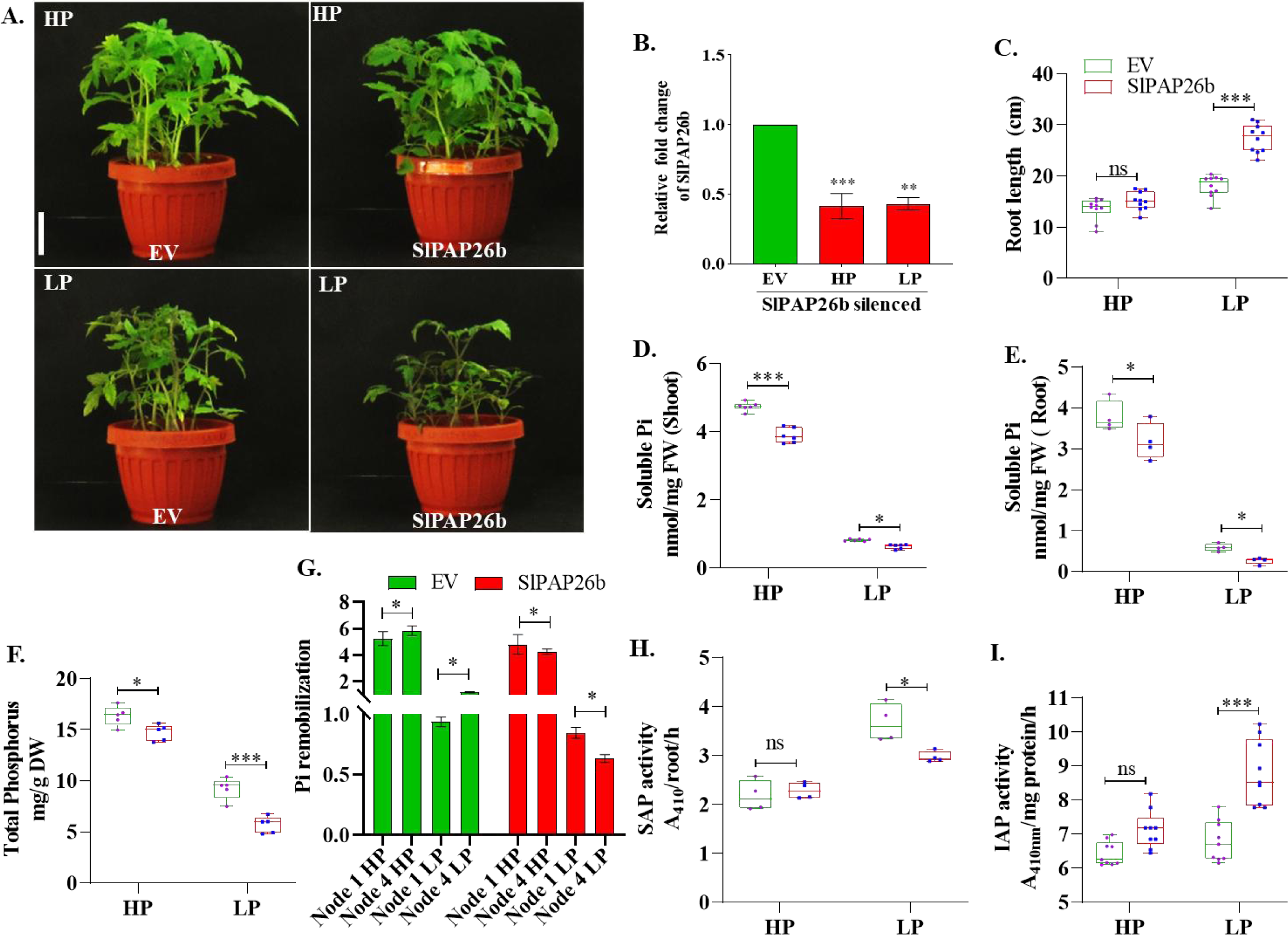
Morpho-physiological and biochemical characterization of *SlPAP26b* silenced tomato seedlings for their P starvation response. **(A)** The phenotype of *SlPAP26b* silenced plants and EV controls under Pi-sufficient (HP) and Pi-deficient (LP) conditions. **(B)** Confirmation of *SlPAP26b* silencing at transcripts level using RT-qPCR **(C)** Primary root length. **(D, E)** Total soluble Pi content in shoot and root tissues. **(F)** Total P content **(G)** Pi remobilization analysis from node 1 to node 4 leaves under HP/LP conditions. **(H, I)** Root-associated total secretory acid phosphatase activity (SAP) and IAP activity. Each asterisk represents the statistically significant differences based on two-way ANOVA tests [(p-value < 0.12 (ns), 0.033 (*), 0.002 (**), 0.001 (***)]. Error bars indicate standard deviation.

On the contrary, the silencing of *SlPAP26b* resulted in stunted seedlings with shorter shoots under Pi-deficient conditions, apparently making them more sensitive to Pi starvation **(Fig. 8A).** The *SlPAP26b* silenced seedlings also exhibited significantly longer primary roots, only under Pi-deficient conditions than their EV controls **(Fig. 8C)**. The silenced seedlings accumulated lower total soluble Pi and total P accumulation in both shoot and root tissues under Pi-sufficient and Pi-deficient conditions, indicating role of this acid phosphatase in Pi uptake **(Fig. 8D-F)**. Because arabidopsis and rice PAP26 genes have been implicated in Pi remobilization from older or senescing leaves to younger leaves and higher transcript levels of *SlPAP26b* were observed in the older senescing leaves of tomato seedlings (**Fig. 5C**), we next investigated the Pi levels in the leaves of different nodes of the silenced and non-silenced plants. This analysis revealed reduced Pi remobilization from node-1 to node-4 leaves in the *SlPAP26b* silenced seedlings compared to their respective EV controls **(Fig. 8G)**. SAP and IAP activities remained unchanged in the *SlPAP26b* silenced seedlings under Pi-sufficient conditions. In contrast, we noticed a 10% reduction in the SAP activity in the root tissue of the *SlPAP26b* seedlings under Pi-deficient conditions **(Fig. 8H**). Contrary to the reduced SAP activity, a concurrent increase in the shoot IAP activity (increased by 22%) was observed in the silenced seedlings over their EV controls under Pi-deficient conditions **(Fig. 8I**). To reconfirm the increased APase activity in the *SlPAP26b* silenced seedlings, we performed in-gel APase assay with total root and shoot tissue proteins of the silenced seedlings and their EV counterparts. More intense bands in both root and shoot tissue gels indicated higher total APase activity in the silenced plants over their EV counterparts under Pi-deficient conditions **(Fig. 9A, B)**. More intensely stained additional bands were observed in the in-gel assays of the shoot tissues of EV and *SlPAP26b* silenced plants under low phosphate conditions. An enrichment of band intensities in the range of 60-75 kDa region indicated activation of PAPs whereas a more intense band at around 100 kDa position suggests that even higher molecular weight acid phosphatases are activated in the shoots of silenced plants than their EV control under Pi deficiency **(Fig. 9A, B)**. These observations also supported the enhanced IAP activity observed earlier in the biochemical assay **(Fig. 8I**, **Fig. 9A, B)**.

**Fig. 9:**
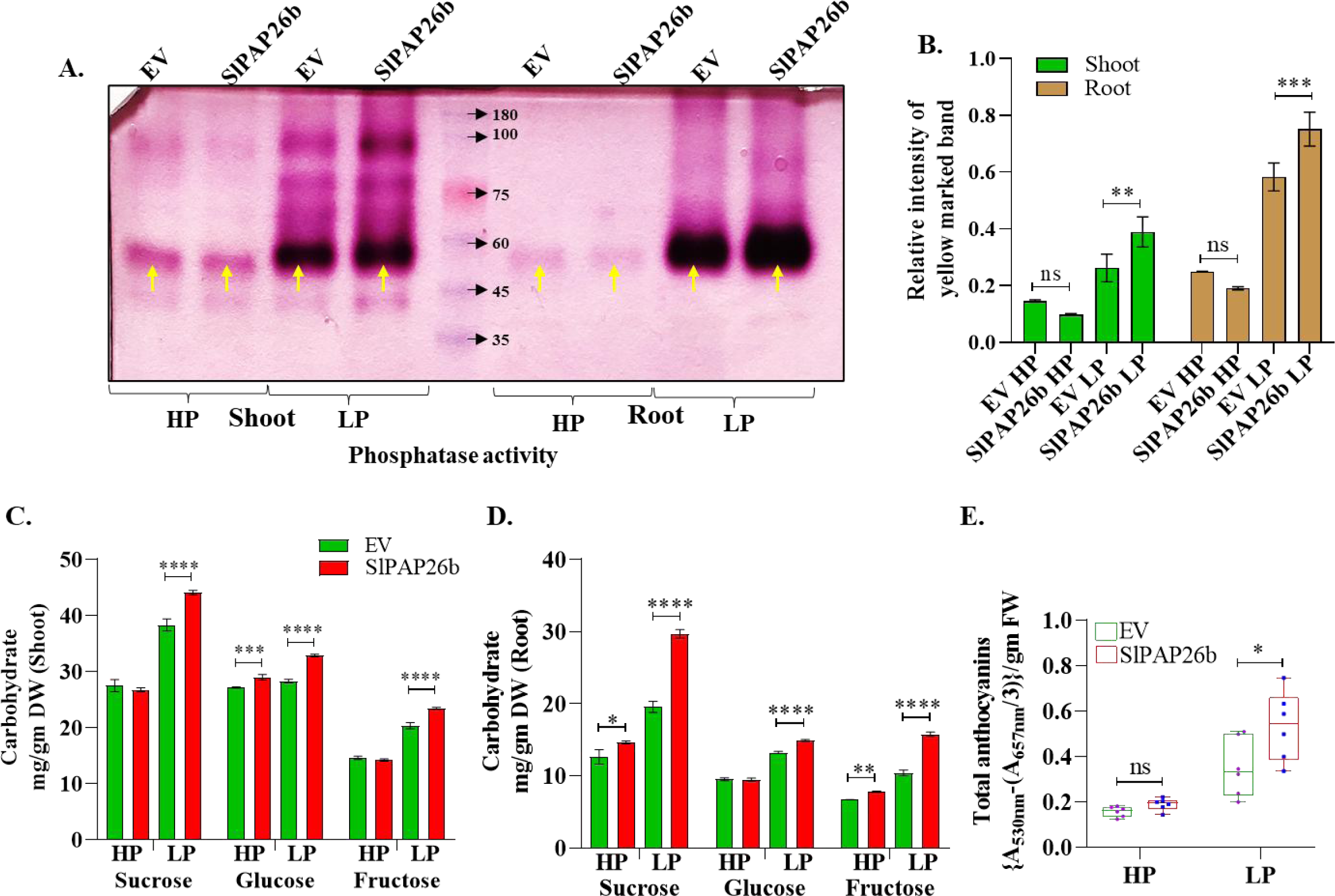
Biochemical characterization of *SlPAP26b* silenced seedlings for APase activity, carbohydrates, and total anthocyanins. **(A)** In-gel assay in *SlPAP26b* silenced and EV plants were performed in root and shoot tissues separately. Each band in the represented gel shows different isoforms of acid phosphatases. **(B)** Relative quantification of APase activity was quantified using yellow marked band by ImageJ software. **(C-D)** Estimation of monosaccharide and disaccharide sugars in silenced and EV seedlings was measured in root and shoot tissues. **(E)** Total anthocyanins content and expression level of its biosynthesis-related genes. Asterisk represent the statistically significant differences based on two-way ANOVA tests [(p-value < 0.12 (ns), 0.033 (*), 0.002 (**), 0.001 (***)]. Error bars indicate standard deviation.

The morphometric analysis and biochemical assays indicated a more severe P starvation response in the *SlPAP26b* silenced seedlings than their counterparts. If SlPAP26b encodes a major APase and its silencing led to accentuated P starvation response, the *SlPAP26b* silenced seedlings are expected to accumulate higher carbohydrate levels along with other P starvation response traits, as described earlier in Fig. 1. As anticipated, enhanced total carbohydrates (sucrose, glucose and fructose), and higher total anthocyanins content supported the invigorated P starvation response in the *SlPAP26b* silenced plants, mainly under Pi-deficient conditions **(Fig. 9E).** A more pronounced induction of anthocyanin biosynthesis genes, such as *SlDFR (Dihydroflavonol 4-reductase)* and *SlF3H (Flavonoid 3′-hydroxylase)* **(Supplementary Fig. Sf6 D)**, or PHT1 transporters, such as *SlPT1* and *SlPT7,* or PSI genes of glycerolipid metabolism pathway, such as *SlGDPD1* (*Glycerophosphodiester phosphodiesterase)*, *SlMGD2* (*Monogalactosyldiacylglycerol synthase2)*, and *SlDGD2* (*digalactosyldiacylglycerol synthase2*) in *SlPAP26b* silenced plants under Pi-deficient conditions supported the biochemical evidence even at the molecular level **(Fig. 8C, 10A)**. Interestingly, the activation of most of these genes was more vigorous in root than shoot tissue of the *SlPAP26b*-silenced seedlings than their EV counterparts **(Fig. 10A)**. To check whether *SlPAP26b*-silencing affected the expression of any vacuolar PHTs, we studied the transcript levels of vacuolar PHTs (*SlPHT5;1*, *SlPHT5;2*, and *SlPHT5;3*) and noticed their accentuated upregulation in the silenced seedlings than their EV counterparts, especially under Pi deprivation **(Fig. 10A)**. Marker PSI genes such as *SlTPS1* and *SlmiR399* also showed stronger induction in their transcripts in the *SlPAP26b* silenced seedlings than EV controls **(Fig. 10A)**.

**Fig. 10:**
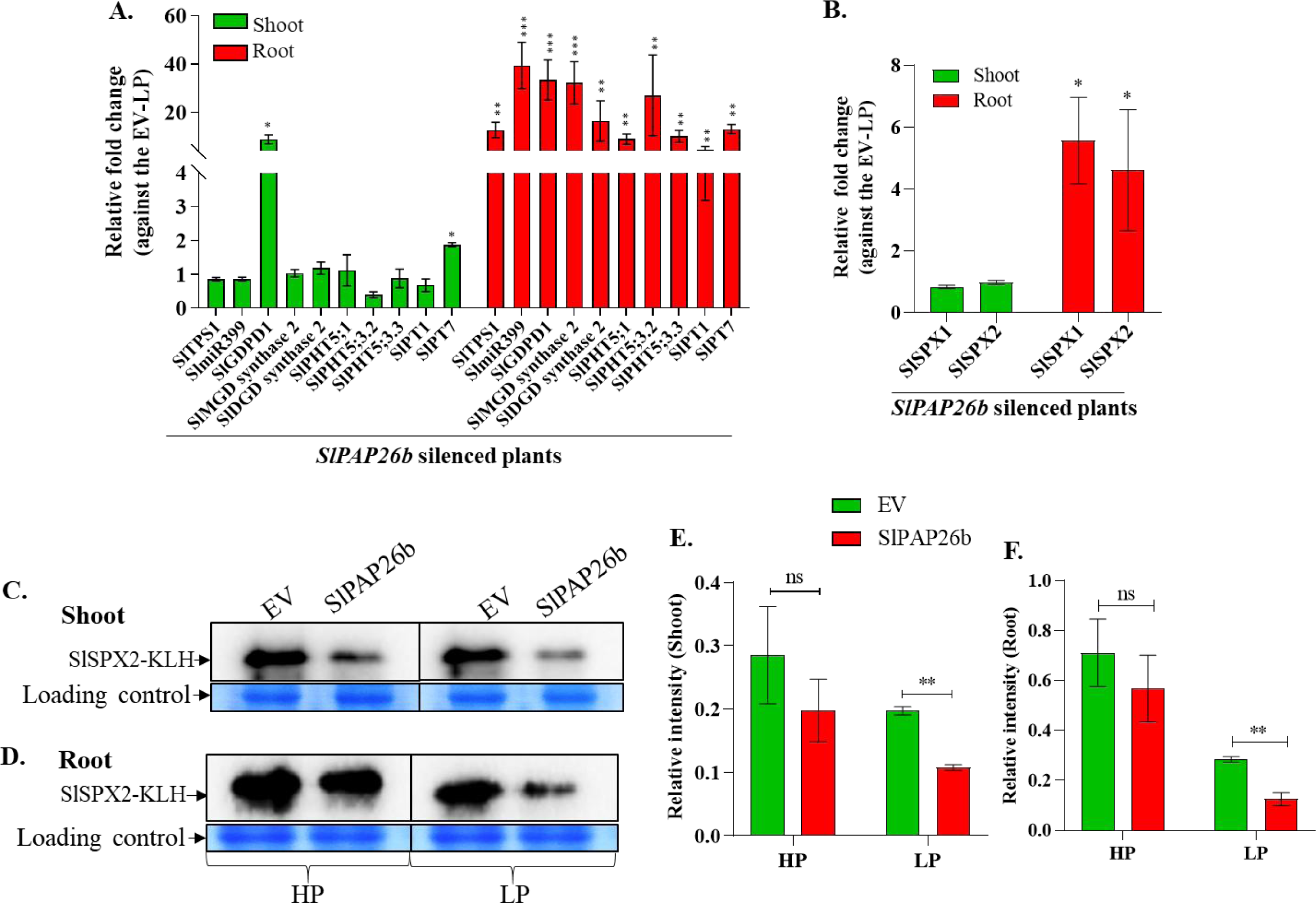
*SlPAP26b* silencing leads to stronger induction of PSI genes in tomato seedlings. **(A)** RT-qPCR based expression profile of PSI genes in *SlPAP26b* silenced root and shoot tissues under Pi deficient (LP) condition. **(B)** Relative mRNA abundance of *SlSPX1* and *SlSPX2* in *SlPAP26b* silenced root and shoot tissues than their EV counterparts under Pi deficient (LP) condition. **(C, D)** Western blot analysis using anti-SlSPX2 antibody and non-native PAGE. Western blot analysis in silenced and EV plants under Pi-sufficient (HP) and Pi-deficient (LP) conditions with SlSPX2-KLH conjugated antibody. **(E, F)** Band intensity from the western blot was quantified using ImageJ software. Asterisk represent the statistically significant differences based on two-way ANOVA tests [(p-value < 0.12 (ns), 0.033 (*), 0.002 (**), 0.001 (***)]. Error bars indicate standard deviation.

Transcriptional regulation of PSI genes is under the tight control of the SPX-PHR1 regulatory module in plants (Srivastava et al., 2021b). If *SlPAP26b*-silenced plants were sensing severe Pi deprivation, then the level of SlSPX proteins is expected to be lower in these seedlings than in their EV controls. To validate the hypothesis, we first checked the transcript levels of PSI *SlSPX1* and *SlSPX2* genes (Singh et al., 2023). As expected, both genes were highly induced at the transcript level in the root tissue of the *SlPAP26b* silenced plants than their EV controls. However, no such change was noticed in shoot tissue under Pi deficient conditions, indicating the more pronounced effect of *SlPAP26b* silencing in roots **(Fig. 10B)**. Next, we performed western blot analysis with total root and shoot proteins using anti-SlSPX2 antibody. Lower accumulation of SlSPX2 protein in *SlPAP26b* silenced seedlings was noticed in the shoot and root tissue, more drastic under Pi-deficient conditions. Altogether, the morphometric analysis, biochemical assays, and molecular results supported the major role of *SlPAP26b* encoded acid phosphatase in the Pi compensatory network in tomato seedlings **(Fig. 10C, D, E, F).**

In contrast to the severe PSR observed in *SlPAP26b* silenced plants, no visible changes in seedlings morphology in *SlPAP17b* and *SlPAP26a* silenced seedlings indicated a limited role of these genes in Pi-homeostasis in tomato (**Supplementary Fig. Sf7, Sf8).** In the literature, a compensatory increase in the transcript levels of other acid phosphatases has been reasoned as one of the main factors for the lack of an apparent P starvation response phenotype in *atpap17* mutant plants (O’Gallagher et al., 2022). To investigate if a similar mechanism also operates in tomato, we examined the transcript levels of selected PAPs in the VIGS-silenced seedlings of these two candidate PAPs. We noticed a higher induction of *SlPAP26b* alongside a modest increase in *SlPAP26a* mRNA levels in the *SlPAP17b* silenced root tissue **(Fig. 11A)**. Similarly, *SlPAP15*, *SlPAP17b,* and *SlPAP26a* were upregulated in *SlPAP26b* silenced seedlings than their EV controls **(Fig. 11B).**

**Fig. 11:**
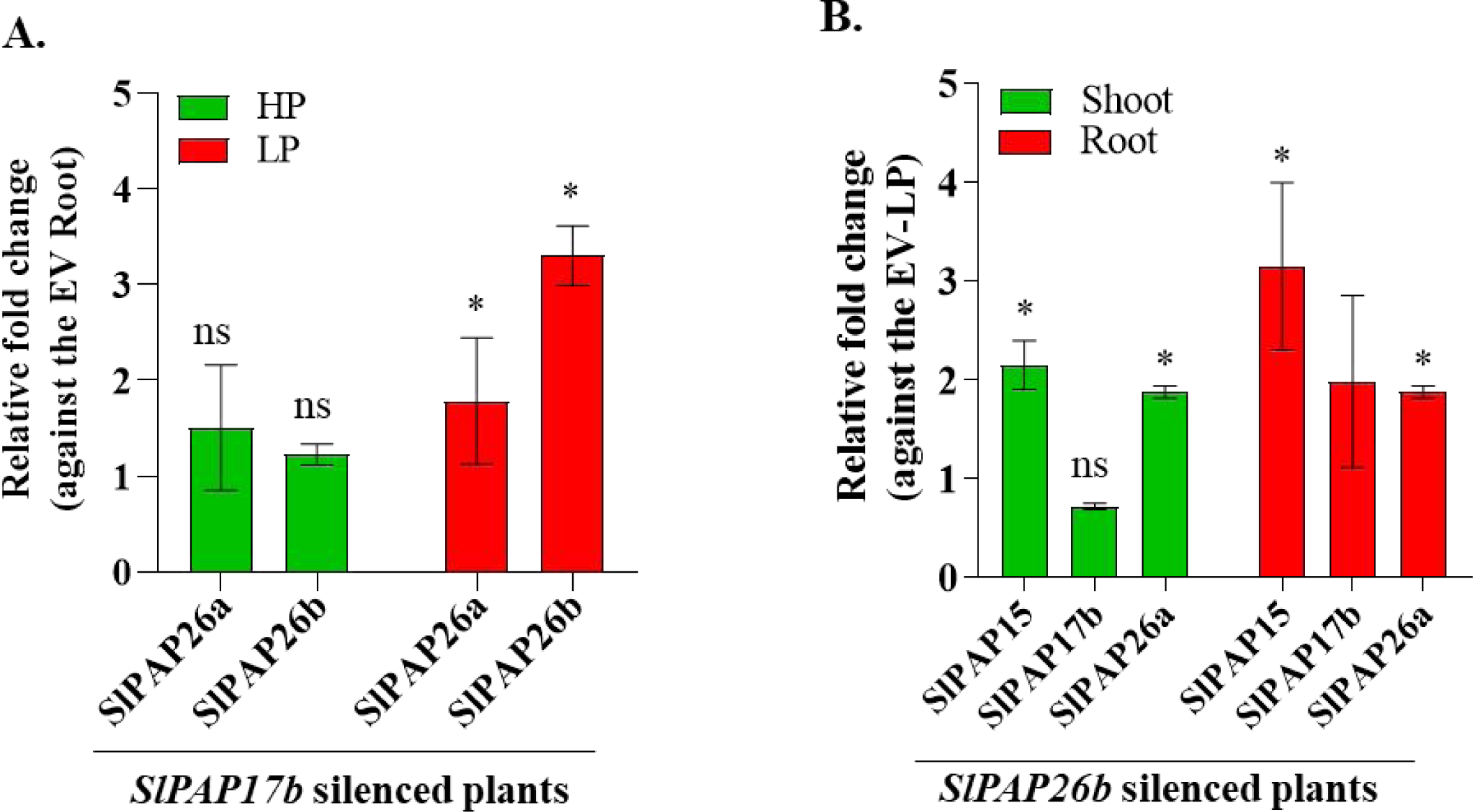
Activation of *SlPAP17b* in *SlPAP26b*-silenced and vice-versa seems to work together in Pi-compensatory networks. **(A)** The expression level of *SlPAP26a* and *SlPAP26b* in *SlPAP17b* silenced roots under Pi-sufficient (HP) and Pi-deficient (LP) conditions. **(B)** The transcript levels of *SlPAP15, SlPAP17b*, and *SlPAP26a* were assessed in the root and shoot of *SlPAP26b* silenced plants under Pi-sufficient (HP) and Pi-deficient (LP) conditions. In both cases, the relative expression of genes was determined against the transcript levels in their respective EV controls. Asterisk represent the statistically significant differences based on two-way ANOVA tests [(p-value <0.12 (ns), 0.033 (*)]. Error bars indicate standard deviation.

### PHR1 homologs fail to bind to *SlPAP26b* promoter in *N. benthamiana*

Due to its PSI nature and major PAP activity, we next investigated whether the well-characterized transcription factors such as *SlPHR1/SlPHL1* (Zhang et al., 2021) can activate the promoter of the *SlPAP26b* gene. Although the in-silico study of the 1.5 kb upstream promoter sequence of the *SlPAP26b* gene did not reveal the presence of a canonical P1BS element, we could identify 10 non-canonical copies of R1BS (RIL-binding site, NAKATNCN) element (Guo et al., 2022). We next tested if the non-canonical P1BS or any other sequence in the 1.5 kb region can serve as the binding site for SlPHR1 or SlPHL1 for its activation under Pi starvation. For this purpose, we performed an in-planta transcriptional reporter activation assay of the *SlPAP26b* promoter (*SlPAP26bpro::GUS*) by SlPHR1 or SlPHL1 in *N. benthamiana* leaves. The promoter of *SlPHT1;2* served as a positive control in the experiment **(Fig. 12A, B)**. While enhanced GUS activity was observed in the case of *SlPHT1;2* promoter, surprisingly, no GUS accumulation was observed in the leaves infiltrated with *SlPAP26b* promoter and either of these transcription factors, suggesting that this gene might not serve as a direct target of by SlPHL1 and SlPHR1 **(Fig. 12A, C)**. To further validate this finding, we checked the transcript levels of *SlPAP26b* in double silenced *SlPHR1+SlPHL1* or *SlSPX1+SlSPX2* VIGS seedlings (Singh et al., 2023). The unaltered transcript levels of *SlPAP26b* in any of the *SlPHR1/SlPHL1* or *SlSPX1/SlSPX2* silenced plants indicated its activation in Pi-deficient tomato seedlings is independent of these two master regulators (**Fig. 12D, E**). In contrast, downregulation of *SlPT1* and *SlPT7,* the known targets of SlPHL1, was observed in the *SlPHR1/SlPHL1* silenced seedlings. The in-planta *SlPAP26bpro::GUS* transactivation assay along with the transcript profile of *SlPAP26b* in *SlPHR1/SlPHL1* or *SlSPX1/SlSPX2* inhibited backgrounds indicate *SlPHR1/SlPHL1* independent upregulation of *SlPAP26b* under Pi deprivation.

**Fig. 12:**
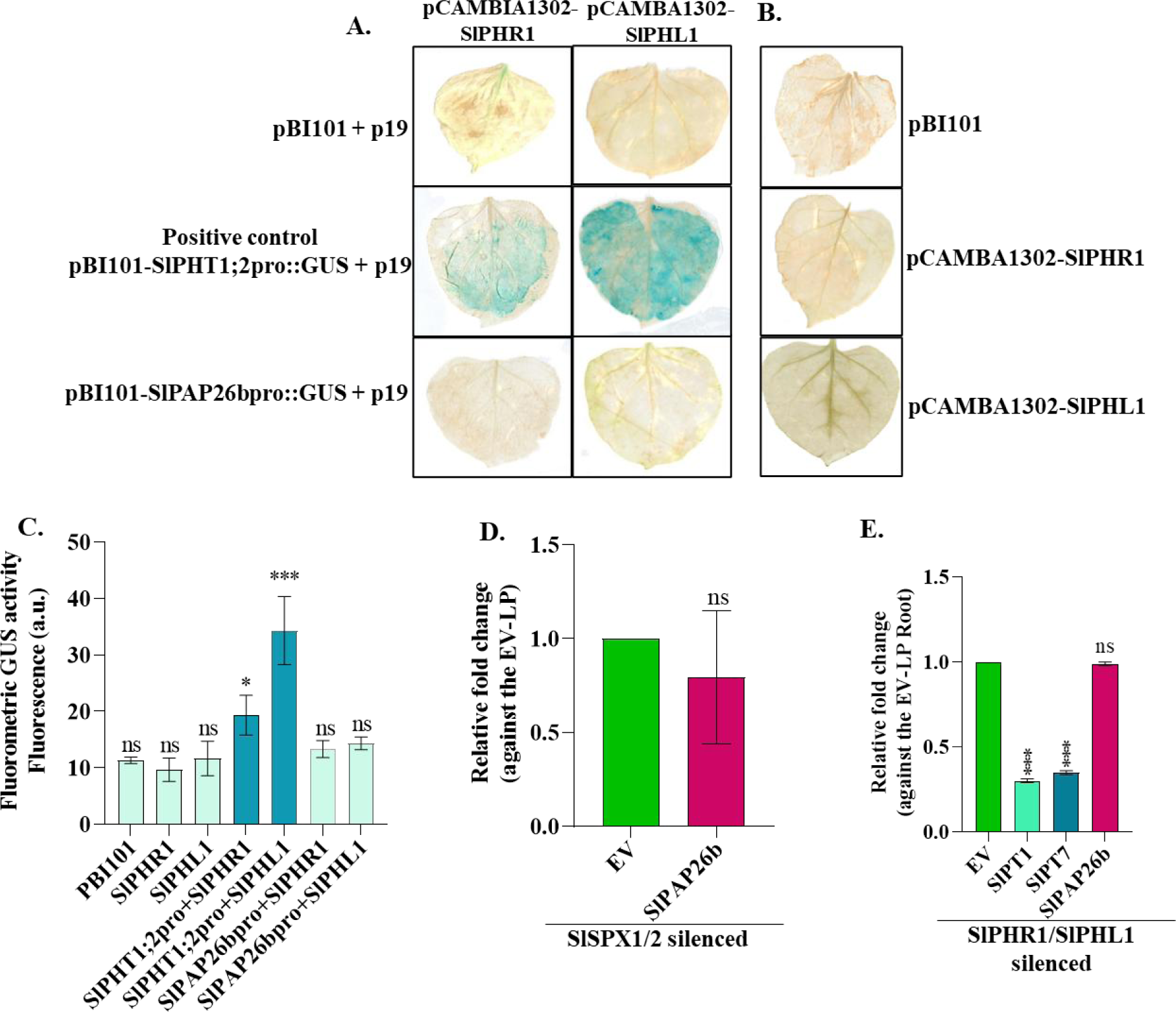
Transcriptional activation assay of *SlPAP26b* promoter by SlPHR1 and SlPHL1. Transient transcriptional activation of *SlPAP26bpro:GUS* promoter via binding of SlPHR1 or SlPHL1. **(A)** GUS assay for *SlPAP26b* promoter activity with SlPHR1 and SlPHL1 homologs. **(B)** Fluorometric GUS estimation by using Fluorescent β-Galactosidase assay (MUG). **(C)** Transcripts level of *SlPAP26b* in *SlSPX1*+*SlSPX2* double silenced seedlings. **(D)** Transcripts level of *SlPAP26b* in *SlPHR1+SlPHL1* double silenced seedlings. *SlPT1* and *SlPT7* acted as positive controls. Asterisk represent the statistically significant differences based on two-way ANOVA tests [(p-value < 0.12 (ns), 0.033 (*), 0.002 (**), 0.001 (***)]. Error bars indicate standard deviation.

## Discussion

Under most natural conditions, Pi is one of the least available nutrients in soils as its concentration is generally low and ranges between 0.65 μM to 2.5 μM (Barber et al., 1963; Karthikeyan et al., 2007). Plants employ an array of well-coordinated morphological, physiological, biochemical, and molecular adaptations to orchestrate their response under chronic phosphate limitation (Plaxton and Tran, 2011). Acid phosphatase activation is an essential aspect of P nutrition and starvation response in plants. Intracellular and secretory acid phosphatases are activated to remobilize Pi from its fixed forms, such as phosphate esters inside and outside plants (Tran et al., 2010). Among all acid phosphatases, purple acid phosphatases form the predominant class of enzymes which contributes to the different aspects of Pi nutrition in plants (Mehra et al., 2017; Ramaiah et al., 2011).

Previous studies have implicated four purple acid phosphatase genes in regulating enhanced PAP activity in tomato (G. Bozzo et al., 2006; Suen et al., 2015). Additionally, *LePS2*, a close homolog of *OsHAD1*, has been characterized as a Pi-responsive gene with APase activity (Baldwin et al., 2001; Baldwin et al., 2008). The activation of several acid phosphatases, including ten purple acid phosphatases *SlPAP1, SlPAP10b, SlPAP12, SlPAP15, SlPAP17b, SlPAP26a*, and *SlPAP26b* in transcriptome and proteome data, and the concurrent enhanced IAP and SAP activity in the Pi-deficient seedlings in the present study reaffirmed the involvement of multiple acid phosphatases, including PAPs, in tomato P starvation response (Bosse & Köck, 1998; G. Bozzo et al., 2002, 2006; G. Bozzo & Plaxton, 2008; Liao et al., 2003; Srivastava et al., 2020; Suen et al., 2015). However, the contribution of most of these PSI acid phosphatases to tomato P starvation response remains unknown.

*AtPAP17* and *AtPAP26* are the main regulators of Pi acquisition and remobilization in Arabidopsis. These two members have dual localization to the cell vacuole and extracellular matrix (O’Gallagher et al., 2022). Thus, both these genes are also secretory in nature (Farhadi et al., 2020; Hurley et al., 2010; Jamali Langeroudi et al., 2023; Robinson et al., 2012). Our search for the homologs of these genes identified two isoforms each for *AtPAP17* (*SlPAP17a* and *SlPAP17b*) and *AtPAP26* (*SlPAP26a* and *SlPAP26b*) in the tomato genome. In silico analysis using PredictProtein predicts the vacuolar/secretory characteristics of both (*SlPAP26a* and *SlPAP26b*) and the secretory/cytoplasmic nature of (*SlPAP17a* and *SlPAP17b*) (Srivastava et al., 2020). Because of the non-PSI nature, highly floral-specific transcript accumulation, and almost identical but weaker expression profile to that of *SlPAP17b*, *SlPAP17a* does not seem to encode a major acid phosphatase in tomato. Moreover, no morphological changes and milder biochemical alterations in the *SlPAP17b-*silenced plants under Pi deprivation indicate a limited role of this gene in the Pi compensation network in tomato seedlings. The limited function of this gene in Pi nutrition could be attributed to its restricted floral-specific induction. Alternatively, the lack of a severe P starvation response phenotype in the *SlPAP17b*-silenced plants could be due to the activation of some other PAPs compensating for its compromised function. This observation is well supported by the *atpap17* mutant seedlings phenotype, where no change in the seedling phenotype of this mutant vis-à-vis wild-type plants has been reported (Jamali Langeroudi et al., 2023). In *atpap17* mutant seedlings, the complementary induction of *AtPAP26* has been reported to minimize Pi starvation response (Farhadi et al., 2020). Upregulation of *SlPAP26a* and *SlPAP26b* transcripts in *SlPAP17b*-silenced tomato roots is in line with earlier observations and suggests that induction of *SlPAP26* isoforms could compensate for the inhibited function of *SlPAP17b* upon its silencing. However, the reduced PUE and total soluble Pi content in *SlPAP17b* silenced seedlings indicate a functional divergence of this gene from its Arabidopsis homolog. Recently, it has been reported that overexpression of *AtPAP17* in soybean improves PUE in transgenic plants. This observation is in line with the altered PUE results presented in the present study (Xu et al., 2022). Overall, the result supports that compensatory acid phosphatases are activated to minimize the impact of the loss of PAP17 orthologs on Pi nutrition in plants.

*AtPAP26* and its rice homolog have been implicated in delayed leaf senescence and Pi remobilization (Robinson et al., 2012; Stigter and Plaxton, 2015). The highly distinct transcript profiles of *SlPAP26a* and *SlPAP26b* during vegetative and reproductive development suggest functional divergence for these two tomato isoforms. The strong and ubiquitous expression profile of *SlPAP26b* in vegetative and floral tissues over *SlPAP26a* and *SlPAP17* isoforms indicates a more profound role of this acid phosphatase throughout plant development. The stronger upregulation of *SlPAP26b,* but not *SlPAP26a*, in the transcriptome and proteome data, also supported that assumption. The stunted shoot growth phenotype of *SlPAP26b* silenced plants, increased anthocyanin content, and longer primary roots under Pi-deprived conditions confirmed the predominant role played by *SlPAP26b* over its tomato paralog in Pi nutrition. Moreover, the unaltered plant phenotype and unchanged soluble Pi content, SAP, and total anthocyanin content in *SlPAP26a* silenced seedlings than their EV controls pointed towards a limited role of this gene in Pi homeostasis in tomato. The kinetics of its activation and recovery experiments indicated a shoot-preferential role of *SlPAP26b* under Pi deprivation. The decreased Pi and total P content and lower Pi remobilization from 2^nd^ to 4^th^ node leaves in *SlPAP26b* silenced Pi-deficient tomato seedlings over their respective EV controls is reminiscent of similar observations reported previously on *atpap26* mutants and in rice (Gao et al.; Hurley et al., 2010; Robinson et al., 2012). Together, these results support a conserved function of PAP26 orthologs in Pi remobilization and acquisition in plants. Stronger activation of the PSI genes such as *SlPT1*, *SlPT7, SlPHT5;1; SlPHT5;3.2/3.3, SlPAP15*, *SlPAP17b,* and *SlPAP26a* in the *SlPAP26b* silenced Pi-deficient seedlings than their respective EV controls indicates exacerbated P starvation response in these plants. The activation of multiple PAPs also explains the mildly reduced SAP activity in roots under Pi deprivation against the anticipated stronger inhibition of APase activity in the *SlPAP26b* silenced seedlings (Farhadi et al., 2020).

Besides a vital role in root growth and expansion, carbohydrates such as sucrose and glucose are known to increase the upregulation of PSI genes in plants (Akash et al., 2021; Franco-Zorrilla et al., 2004; Hammond and White, 2008, 2011; Karthikeyan et al., 2007; Khurana et al., 2021; Müller et al., 2007; Yang et al., 2020; Zhu et al., 2005). The increased levels of several sugar metabolites in Pi-deficient seedlings, longer primary root length in (-P+S) seedlings, and higher APase activity in both Pi-sufficient and Pi-deficient seedlings with sucrose at 8-D/15-D time points in the present study are in line with the previous reports (Karthikeyan et al., 2007; Zakhleniuk et al., 2001). The significantly enhanced levels of sucrose, D-glucose, and D-fructose in shoot and root tissues of *SlPAP26b*-silenced Pi-deficient tomato seedlings than their respective EV controls indicates that higher carbohydrates levels could be one of the primary reasons for the exacerbated P starvation response, including longer roots, enhanced APase activity and elevated upregulation of PSI genes observed in the silenced plants (Hammond and White, 2008; Karthikeyan et al., 2007; Lei et al., 2011; Liu et al., 2005; Nacry et al., 2005; Zakhleniuk et al., 2001). The stronger activation of PSI genes in the root tissue of *SlPAP26b-*silenced Pi-deficient seedlings over their respective EV control is also validated by the significantly decreased levels of SlSPX2 protein, a root-preferential gene (Singh et al., 2023). The stronger activation of PSI genes in the root than the shoot, even though SlSPX2 levels was significantly reduced in both tissues, of *SlPAP26b* silenced plants is intriguing and could be either related to the root preferential nature of *SlSPX2* or enhanced sucrose remobilization from shoot to root under Pi deficiency. Alternatively, it could be due to the root preferential nature of *SlPHL1*, the main PHR1 homolog anticipated to become active on PSI genes after the degradation of SlSPX2 in tomato roots under Pi deficiency (Zhang et al., 2021; **Supplementary Fig. Sf9**).

Transcription of PSI genes is tightly controlled by MYB transcription factors, PHR1, and its homologs in plants. Under Pi-sufficient conditions, SPX proteins (the negative regulators of P starvation response) physically interact with PHR1 homologs and regulate their transcription factor activity to control the expression of PSI genes (Rubio et al., 2001; Wang et al., 2014). Due to the exclusive PSI nature of *SlPAP26b* and severe P starvation response phenotype observed upon its silencing in tomato seedlings, we next investigated whether the SlSPX-SlPHR module directly regulates *SlPAP26b* transcript levels. The unchanged transcript levels of this gene in *SlSPX1/SlSPX2* or *SlPHR1/SlPHL1* silenced plants pointed to the limited role of SlPHR1 and SlPHL1 in the direct activation of this gene upon Pi deprivation. Further, no GUS activity in the in-planta transactivation *SlPAP26bpro::GUS* transient assays indicates SlPHR1 or SlPHL1 independent activation of this gene in Pi-deficient seedlings. The absence of the P1BS element in the promoter region of its rice homolog, *OsPAP26*, also supports PHR1-independent regulation of PAP26 homologs in plants (Zhang et al., 2011). Recently, Liao et al. (2022) reported the presence of 17 PHR1 homologs in tomato (Liao et al., 2022). Additionally, RLI1a (REGULATOR OF LEAF INCLINATION1), another MYB class transcription factor, has also been identified to activate the expression of PSI genes by binding to the R1BS (NAKATNCN) element (Guo et al., 2022; Ruan et al., 2018). In light of these recent reports and the fact that *SlPAP26b* is exclusively activated only under Pi deprivation, we cannot rule out the involvement of another tomato PHR1 homolog or an OsRLIa homolog in regulating the expression of this gene.

In summary, the enhanced APase activity in Pi-deficient tomato seedlings is contributed by the activation of many acid phosphatases, including a major contribution coming from purple acid phosphatases. Elevated levels of sugars seem to affect APase activity by enhancing the transcription of acid phosphatase genes and also by promoting sugar-phosphates metabolites in tomato seedlings, thus decreasing the effective concentration of cellular Pi especially under its limitation. Unaltered transcript levels of *SlPAP26a* show that the expression of not all PAPs responds positively to the sucrose treatment. The results presented here support the earlier observations that the activation of other compensatory PAPs may help minimize the impact of the loss of function of a purple acid phosphatase during Pi nutrition in plants. The detailed characterization of three candidate PAPs and only visibly altered phenotype in *SlPAP26b* silenced seedlings indicate that *SlPAP26b* encodes a major purple acid phosphatase in tomato. Its ubiquitously stronger expression, mainly in senescing leaves than younger leaves, and activation upon *SlPAP17b* silencing in the *SlPAP26b* silenced plants could be one of the reasons for the mitigated PSR response observed in those plants. The overall lower P content and compromised Pi remobilization in the *SlPAP26b* silenced plants than their EV control further supports a major role of *SlPAP26b* in Pi remobilization in tomato seedlings. Although specifically induced upon Pi starvation, its unchanged transcript levels in either SPXs or PHRs silenced plants and the lack of a P1BS element in its 1.5 kb promoter region is intriguing and indicates that *SlPAP26b* transcription is regulated independent of SlPHR1 and SlPHL1. Although we failed in inducing recombinant SlPAP26b protein expression in *E. coli* (data not shown), it will be interesting to find out the substrates for this major acid phosphatase and the factors responsible for its specific activation upon Pi-deficiency, including the newly identified tomato PHR1 homologs.

## Acknowledgment

Science and Engineering Research Board, Govt. of India (grant no. CRG/2018/001033) supported this work. Akash and RS acknowledge the Council of Scientific and Industrial Research, Govt. of India, for JRF and SRF fellowships. AR acknowledge UOH-BBL and UOH-IOE incentive for fellowship. The authors also acknowledge the institutional contributions from the Department of Science and Technology (DST), Government of India, Funds for Infrastructure in Science and Technology (FIST), Level II, the University Grants Commission supported Special Assistance Programme (UGC-SAP-DRS-II) to the Department of Plant Sciences, University of Hyderabad (UoH) and to the University of Hyderabad-IoE by MHRD (F11/9/2019-U3(A) and DBT-support to the School of Life Sciences under BUILDER program.

## Author contributions

RK conceived the project and designed the experiments. RK, Akash, RS, AR, KS, MC and PK conducted the experiments. Akash, RS, KS, and MC analyzed the data. AR, MC, and PK designed and analyzed the proteomics experiment. RK, Akash, and RS wrote the paper. All authors read and approved the final manuscript.

## Data availability

All data supporting the findings of this study are available within the paper and its supplementary materials published online. RNA-seq data that support the findings of this study have been deposited in the NCBI SRA database under record SUB8104515.

## Supplementary figure legends

**Supplementary Fig. Sf1:** Morpho-physio analysis of 8-D/15-D old seedlings grown under HP/LP condition. **(A, B, C, D)** Root length, lateral root numbers, shoot length, and total anthocyanins content upon 8-D/15-D post-starved seedlings. Asterisk represent the statistically significant differences based on two-way ANOVA tests [(p-value < 0.12 (ns), 0.033 (*), 0.002 (**), 0.001 (***)]. Error bars indicate standard deviation.

**Supplementary Fig. Sf2:** Validation of RNA-seq results by RT-qPCR of the 17 genes that were used in this analysis. The fold-change values obtained for each gene in the two data sets were plotted against each other. **(A, B)** 8-D and 15-D, respectively, under low phosphate conditions. *SlGAPDH* was used as an internal control.

**Supplementary Fig. Sf3: (A)** Heatmap showing levels of the detected metabolites in HP- and LP-grown seedlings. **(B, C)** 3-D and 2-D Principal component analysis of the HP- and LP-grown seedlings (n = 5). The PCA was plotted using the MetaboAnalyst 4.0. **(D, E).** Heatmap showing levels of the detected metabolites in HP- and LP-grown seedlings. Both rows and columns were selected to generate a heatmap in Morpheus (https://software.broadinstitute.org/morpheus/) with hierarchical clustering, euclidean metric, and average linkage method.

**Supplementary Fig. Sf4:** Morpho-physio analysis of 8-D/15-D old seedlings grown under HP/LP condition upon sucrose supplementation. **(A, B, C, D, E, F)** Primary root length, lateral root density, and total anthocyanins content upon 8-D/15-D Pi starved seedlings with/without sucrose supplementation. Asterisk represent the statistically significant differences based on two-way ANOVA tests [(p-value < 0.12 (ns), 0.033 (*), 0.002 (**), 0.001 (***)]. Error bars indicate standard deviation.

**Supplementary Fig. Sf5: (A, B, C, D)** Phylogenetic analysis and multiple sequence alignment of *SlPAP17a*, *SlPAP17b*, *SlPAP26a,* and *SlPAP26b* genes was constructed using PhyloGenes in the SGN tool. **(E, F)** Expression analysis of *SlPAP17a*, *SlPAP17b*, *SlPAP26a,* and *SlPAP26b* genes at different development stages using multi-Plant eFP Browser 2.0 with online expression data.

**Supplementary Fig. Sf6: (A, B)** The photobleached leaf phenotype observed in most of the infiltrated plants (≥80%) confirmed highly effective gene-silencing of *SlPDS*. **(C)** Relative fold change of anthocyanin biosynthesis pathway genes in *SlPAP26b* LP silenced seedlings with respect to EV LP control seedlings. The asterisk represents the statistically significant differences based on unpaired student t-test [(p-value < 0.033 (*)]. Error bars indicate standard deviation.

**Supplementary Fig. Sf7: (A)** Characterization of *SlPAP26a* using VIGS. **(B)** Confirmation of silencing of *SlPAP26a* using RT-qPCR. **(C)** Total soluble Pi content of the *SlPAP26a* silenced seedlings in HP and LP conditions. **(D)** The total P content of the *SlPAP26a* silenced seedlings in HP and LP conditions. **(E)** Secretory APase activity in the *SlPAP26a* silenced seedlings. **(F)** Total anthocyanins contents in the silenced and unsilenced seedlings. Empty vector infiltrated seedlings served as control. Asterisk represent the statistically significant differences based on two-way ANOVA tests (p-value < 0.12 (ns), 0.033 (*), 0.01 (**). Error bars indicate standard deviation.**Supplementary Fig. Sf8:** Characterization of *SlPAP17b* using VIGS. **(A)** Phenotype of silenced plants. **(B)** Confirmation of silencing of *SlPAP17b* using RT-qPCR. **(C)** Total soluble Pi content of the *SlPAP17b* silenced seedlings in HP and LP conditions. **(D)** The total P content of the *SlPAP17b* silenced seedlings in HP and LP conditions. **(E)** Secretory APase activity in the *SlPAP17b* silenced seedlings. **(F)** Phosphate use efficiency *SlPAP17b* silenced seedlings in HP and LP conditions. **(G)** Total anthocyanins content in the silenced and unsilenced seedlings. Empty vector infiltrated seedlings served as control. Asterisk represent the statistically significant differences based on two-way ANOVA tests [(p-value < 0.12 (ns), 0.033 (*), 0.002 (**)]. Error bars indicate standard deviation.

